# Tuning T-cell immunological synapse by modular DNA-Nanobody engagers for precision immunotherapy

**DOI:** 10.64898/2026.07.22.740045

**Authors:** Juliette Prothon, Gaurav Verma, Beatriz Díaz-Bello, Martine Biarnes-Pelicot, Florian Dupuy, Charlie Gosse, Gaetan Bellot, Kheya Sengupta, Laurent Limozin, Patrick Chames

**Author notes:** Equal contribution (Juliette Prothon, Gaurav Verma). Equal contribution (Laurent Limozin, Patrick Chames).

## Abstract

Bispecific T-cell engagers (TCEs) are a promising class of cancer immunotherapies, but their clinical use is limited by toxicity and insufficient specificity. Tuning the T cell–tumor interface through engager architecture may address these drawbacks. To this end, we engineered hybrid constructs composed of two nanobodies targeting CD3 and the model tumor antigen HER2, respectively, connected by rigid DNA linkers of variable length. Using cytotoxicity assays and hybrid biophysical platforms, we demonstrate a linker-length dependence of cell spreading on antigen, target killing and cytokine release, revealing a functional decoupling between killing and cytokine secretion, and implicating the glycocalyx as a key player. Through the addition of EGFR targeting, we also generate trispecific constructs implementing an “OR-gate” logic to address tumor heterogeneity and reduce resistance due to antigen loss. Overall, these versatile constructs show great therapeutic promise, and at the same time serve as platforms to test hypotheses on biophysical mechanisms.

## Introduction

The advent of immunotherapies, particularly T-Cell Engagers (TCEs), has revolutionized oncology by restoring the immune system’s ability to eradicate tumors^1^. TCEs act as molecular bridges, forcing physical proximity between T cells and cancer cells. This co-localization induces the formation of an artificial immunological synapse (IS), a critical interface that facilitates optimal intercellular communication, triggers intracellular signaling pathways, and activates cytotoxic functions involving the secretion of lytic granules.^2^

Unfortunately, in clinical settings, this potent activation is often intrinsically linked to substantial toxicity. TCEs therapies are frequently associated with severe immune hyperactivation, known as Cytokine Release Syndrome. This phenomenon is characterized by a massive release of pro-inflammatory cytokines (TNF-α, IFN-γ, IL-6) that can lead to fatal multi-organ failure, significantly narrowing the therapeutic window of these agents^3^. To mitigate this risk, recent strategies have explored the use of TCEs with reduced affinity^4^. While this approach limits immune overstimulation, it often results in compromised anti-tumor efficacy, necessitating significantly higher doses to achieve a clinical response^5^. Consequently, it is imperative to identify alternative strategies capable of decoupling more efficiently cytotoxic efficacy from systemic toxicity.

Beyond biochemical affinity, cytotoxic efficacy is governed by the architecture of the immunological synapse (IS), which emerges as a major determinant of lymphocyte activation. The formation of the IS is governed by strict biophysical constraints, where molecular geometry and mechanical forces play a predominant role^2^. The kinetic-segregation model notably highlights the importance of the inter-membrane distance: the formation of close contacts favors the spatial exclusion of CD45 (a bulky inhibitory transmembrane glycoprotein phosphatase) from the contact zone^6^. This exclusion is essential to permit the phosphorylation by the kinase Lck of Immunoreceptor Tyrosine-based Activation Motifs (ITAMs) within the T-cell receptor (TCR) complex, thereby initiating the signaling cascade^7^. Thus, the precise modulation of synaptic topography via structural modifications of TCEs represents a promising, yet underexploited, avenue to fine-tune the immune response^7^.

A distinct but equally critical challenge lies in the fact that the clinical efficacy of bispecific TCEs is often compromised by antigenic heterogeneity, both inter- and intra-patient, and by the emergence of secondary resistance driven by antigen loss. To circumvent these escape mechanisms, an interesting strategy consists in the development of trispecific T-cell engagers capable of simultaneously targeting two distinct tumor antigens in addition to CD3, for instance CD19 and CD22 or EGFR and EpCAM ^8,9^. Operating via an “OR-gate” logic, this system triggers T-cell activation upon recognition of at least one of the target antigens, thereby conferring a dual therapeutic advantage. This approach prevents tumor escape by ensuring lysis even if one antigen is lost under selective pressure, and can also exploite the concomitant engagement of both arms by increasing avidity and synapse stability on cells coexpressing both targets.

The rational design of conventional TCE formats is hindered by structural limitations inherent to their nature, relying on the engineering of full-length antibodies or their fragments. While these constructs provide useful insight about the impact of engager flexibility and geometry^10,11^, they suffer from limited fine control over the conformation of the TCE, and a lack of modularity, which complicates rapid adaptation to new molecular targets. Addressing these technological hurdles requires the development of versatile and modular platforms capable of integrating these biophysical parameters to design the next generation of immunotherapies. Thus, recent works have combined artificial or single domain antibodies to produce modular multivalent constructs^12,13^. Alternatively, DNA were coupled to single chain Fv fragments (scFv)^14^, DNA aptamers^15^ or via DNA origamis^16^. However, balancing the geometrical control of binder arrangement and the overall size of the construct, so as to match the immune synapse, remains difficult when using these frameworks.

To overcome these challenges, this work introduces hybrid DNA/protein TCEs we name DNanobodies (DNabs). These constructs synergize the structural rigidity, programmable positioning, and dimensional precision of double-stranded DNA (dsDNA) with the small size, specificity, and modularity of camelid-derived single-domain antibodies (nanobodies or VHH)^17^. DNabs are designed to orchestrate the simultaneous engagement of immune and tumor cells by recognizing specific surface markers. Two architectures were developed to probe distinct mechanistic hypotheses. First, we designed bispecific constructs in which anti-HER2 and anti-CD3 nanobodies are linked via a series of dsDNA spacers. By enabling the precise modulation of the linker length, this format allows a systematic assessment of how inter-membrane distance influences the immunological synapse and functional potency. We also propose a trispecific format incorporating anti-CD3, anti-HER2 and anti-EGFR nanobodies to provide an ‘OR-gate’ Boolean logic and evaluate the contribution of multispecificity to immune recruitment and cytotoxicity induction. By combining molecular, cell-surface and cell-cell interaction assays, this work offers a dynamic evaluation of DNab function at the synapse.

## Results

### Design and production of DNabs

The inter-membrane space between T cells and target cells is a critical parameter for the formation and efficiency of the immunological synapse, directly influencing TCR signaling, T cell activation and cytokine release. Our aim was to design a new generation of bispecific T cell engagers based on DNA-nanobody hybrids combining anti-HER2 and anti-CD3 nanobodies, i.e. small single domain antibodies (15 kDa) derived from camelids, fused via a rigid double strand DNA linker of well-defined length to precisely study the effect of the size of the engagers on their efficacy. Four formats were generated (**Fig. 1a**). Three formats carried nanobodies at both ends of a double strand DNA of different lengths (20, 30 and 40 base pairs (bp)), thus making it possible to control the relative distance between the anti-HER2 and anti-CD3 Nbs, each bp corresponding to 0.34 nm. This resulted in estimated inter-epitope distances of approximately 20, 24, and 27 nm, respectively (**Supplementary Fig. 1**). An additional construct, named 30cis, was designed to position two Nbs on the same side of the DNA double helix, thereby minimizing the effective linker length and yielding an inter-epitope distance of approximately 16 nm (**Supplementary Fig. 1**). Control molecules integrating an irrelevant Nb (Nef19) were also synthesized to rule out artifacts related to DNA structure or non-specific effects (**Fig. 1a**). All DNabs were produced according to a uniform assembly procedure and subsequently purified by size-exclusion chromatography (**Extended Data Fig. 1** and **2a**). Their purity was verified by denaturing SDS-PAGE and agarose gel electrophoresis, and only the pure fractions were retained for further investigations (**Fig. 1b** and **Extended Data Fig. 2b-c**).

**Fig. 1:**
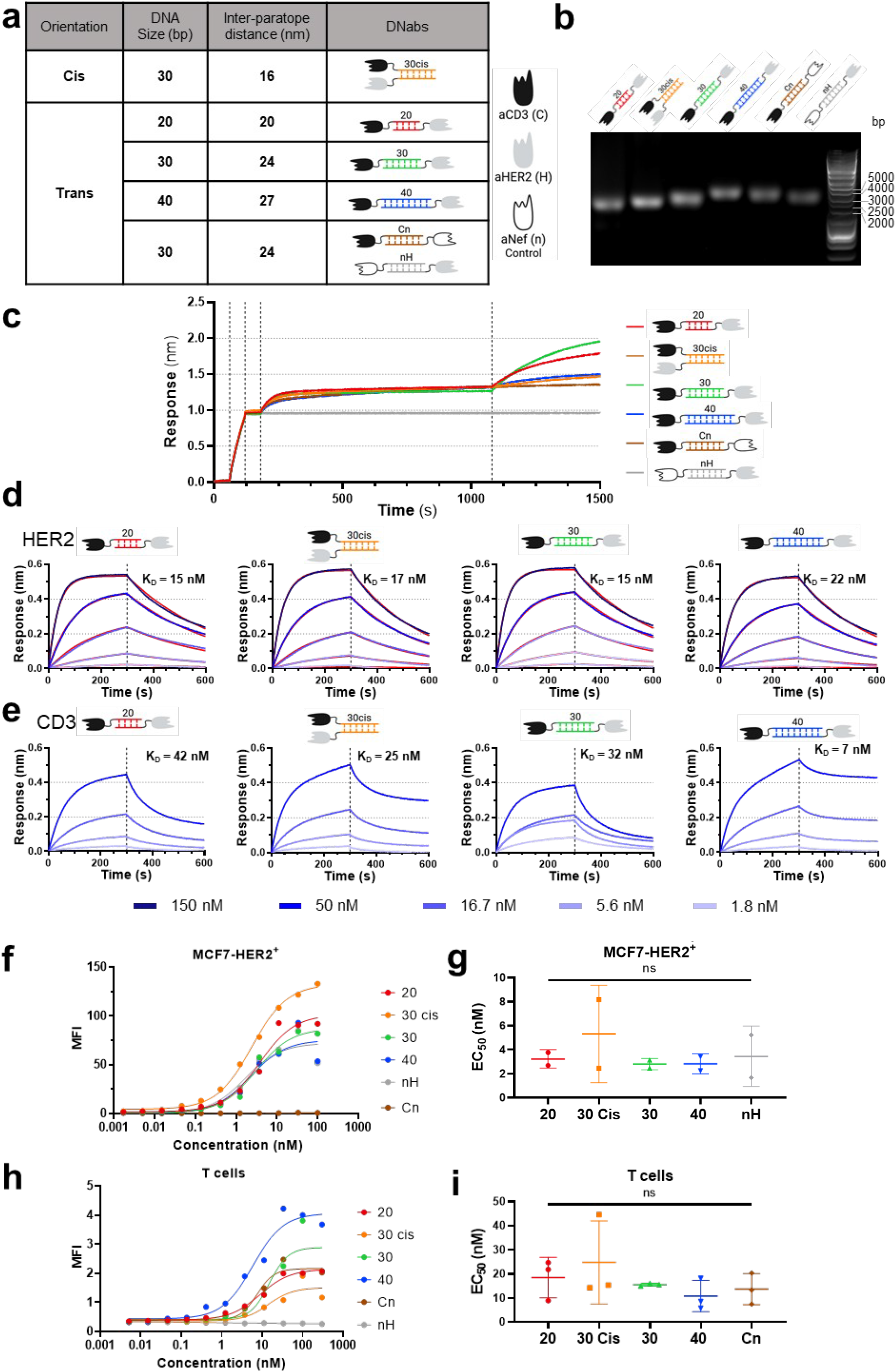
Design and characterization of bispecific anti-HER2 x CD3 DNabs. **a,** Schematic representation of DNabs. The indicated lengths correspond calculated by considering the maximal distance of a fully extended DNab conformation (Supplementary Fig. 1). **b,** Assessment of the construct purity by 1% agarose gel electrophoresis. **c,** Assessment of bispecific binding by Bio-Layer Interferometry (BLI) (typical dataset from n = 3 experiments). Streptavidin biosensors loaded with biotinylated recombinant CD3εδ heterodimer were sequentially exposed to 50 nM of the indicated DNabs (20, 30, 30cis, or 40) or controls (Cn, nH), followed by a secondary association step using recombinant HER2-Fc (10 µg/mL) in the presence of the same DNabs (50 nM). **d,** BLI monitoring of DNabs binding to immobilized biotinylated HER2 nM (typical dataset from n = 3 experiments displayed in Extended Data Fig. 4). Blue curves represent experimental measurements at 4 concentrations (ranging from 1.56 to 150 nM), while red curves correspond to the 1:1 global fit model. **e,** BLI monitoring of DNab binding to immobilized biotinylated CD3εδ. Blue curves represent experimental measurements at 4 concentrations ranging from 1.56 to 50 nM (typical dataset from n = 3 experiments displayed in Extended Data Fig. 5). **f,** Apparent affinity (determined as the half-maximal effective concentration, EC50) measurement by flow cytometry on MCF-7-HER2^+^ cells (typical dataset from n = 2 experiments. **g,** Summary of EC_50_ values obtained from independent flow cytometry experiments on MCF-7-HER2^+^ cells (one-way ANOVA, p = 0.89). Data are presented as mean ± SD, *n* = 2. **h,** EC_50_ measurement by flow cytometry on primary T cells isolated from healthy donors (typical dataset from n = 3 experiments). **i,** Summary of EC_50_ values obtained from independent flow cytometry experiments on T cells (one-way ANOVA, p = 0.79). Data are presented as mean ± SD, *n* = 3. *Abbreviations: aHER2: nanobody anti-HER2; aCD3: nanobody anti-CD3; bp: base pairs; EGFR: Epidermal Growth Factor Receptor; HER2: Human Epidermal growth factor Receptor 2; MFI: Median Fluorescence Intensity*

### Validation of bispecific binding and affinity characterization

The capacity of DNabs to simultaneously engage both targets was initially validated using a two step bio-layer interferometry (BLI) assay. The analysis confirmed the concomitant binding of DNabs to both CD3 and HER2 antigens. As expected, the monospecific controls did not exhibited the same double association profile (**Fig. 1c**). Subsequently, binding kinetics were characterized. All constructs (20, 30cis, 30, and 40) exhibited comparable affinities against immobilized HER2, with respective K_D_ values of 15 ± 3, 17 ± 8, 15 ± 3, and 22 ± 8 nM ( **Fig. 1d, Extended data Figs. 3 and 4**). No statistically significant differences were observed (one-way ANOVA, p = 0.6340) (**Extended Data Fig. 3**). BLI measurements performed with immobilized CD3εδ exhibited complex binding profiles that deviated from simple 1:1 kinetics (**Fig. 1e**). Consequently, K_D_ were determined using steady-state analysis (Extended data **Fig. 5**) Unexpectedly, DNab 40 exhibited a slightly higher affinity compared to the other three formats (p < 0.05; one-way ANOVA followed by Tukey’s multiple comparisons test). The calculated K_D_ were 7 ± 2 nM for DNab 40 compared to 42 ± 9 nM (20), 25 ± 11 (30cis), and 32 ± 15 (30) (Extended data **Fig. 3**).

To validate antigen recognition within their native context, the binding of DNabs was assessed by flow cytometry. Binding experiments performed on MCF-7-HER2^+^ revealed similar apparent affinities across all formats, in the nanomolar range (**Fig. 1f-g, Supplementary Table 1**). The interaction with CD3 was measured on activated primary human T cells and show no significant difference between formats (**Fig. 1h-i**, and **Supplementary Table 1**).

### T Cell engagement on antigen surface depends on DNab length and concentration

The capacity of DNabs to mediate cell engagement on antigen surfaces was studied using brightfield and Reflection Interference Contrast Microscopy (RICM)^18^. Primary human T cells were deposited on an HER2-coated surface in the presence of DNabs at fixed concentration (**Fig 2a**). As cells sedimented, they were detected using brightfield and RICM (**Fig. 2b**), then segmented and tracked using Celldetective software^19^. A hovering phase (after detection but before engagement on the surface) was observed (**Fig 2b, Supplementary Video 1)**. Cell engagement was initiated at spreading onset time t_0_, determined when the cell contact area, as measured in RICM by the surface area of dark pixels, exceeded a threshold of 8 µm² (**Fig 2b**), and was followed by the spreading of cell on the surface. The distribution of hovering durations before engagement was represented as an engagement survival curve (representative curves on **Fig 2c**), whose slope represents the rate of engagement. At a concentration of DNab of 6 nM, the rate of engagement, measured for each DNab (**Fig 2d, Supplementary Video 1)**), exhibited an increase as a function of DNab length. At higher concentration (16 nM), the engagement rate was increased for shorter DNabs (except for 30cis) and the length dependence was partially abrogated (compare **Fig. 2d** and **2e**. See Extended data **Fig. 6a**). All these trends were also evidenced when representing the fraction of detected cells that engaged within their first 10 min of hovering (Extended data **Fig. 6b**). Upon engagement, the cell spreading area was measured as a function of time (inset of **Fig 2f**) and the maximum it reached is determined. The maximal area increased with DNab length (**Fig. 2g, Supplementary Video 1**), a trend that was confirmed at 16 nM, with only marginal gain in spreading (compare **Fig 2g** and **2h, Extended Data Fig. 6d**).

**Fig. 2:**
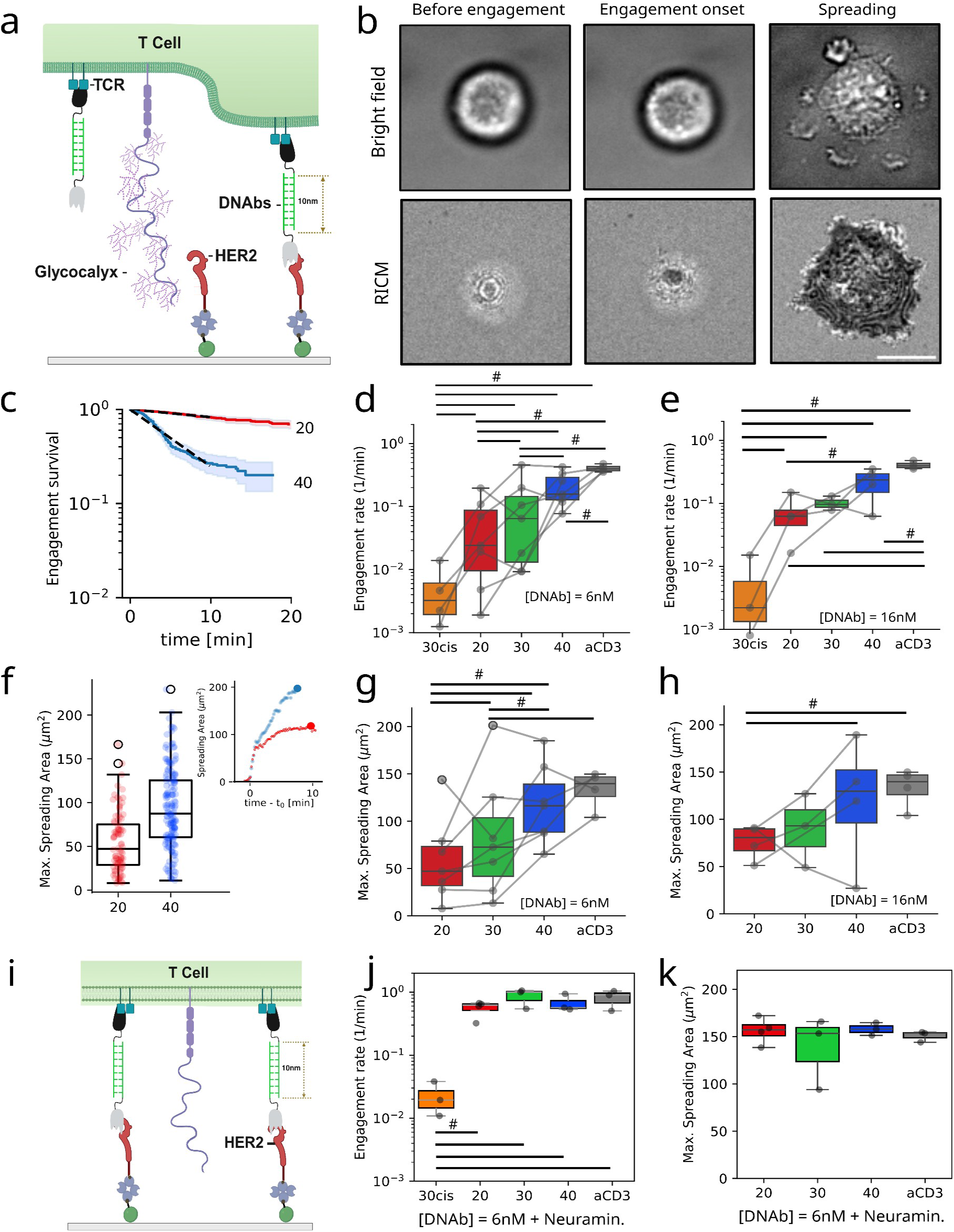
T-Cell-spreading on antigen surface is controlled by DNab format and cell glycocalyx. **a,** Schematics of T-cell interaction with an antigen bearing surface in presence of DNab. **b**, Bright field and RICM micrographs of cell prior to engagement on the surface, at engagement onset or during spreading. Scale bar: 8 µm. **c**, Representative survival curves of the duration between cell detection and engagement on the surface for DNabs 20 and 40. The slope of the dashed line defines the engagement rate. **d,** Engagement rate for all DNab formats (concentration 6 nM, anti-CD3 as positive control). Each point represents one experiment (20-100 cells per experiment). **e,** Engagement rate for all DNab formats (concentration 16 nM). **f**, Example of Maximal Spreading Area for one experiment and 2 DNabs. Each point represent one cell. Inset: Representative spreading area curves as a function of time after engagement at onset time t_0_. **g,** Maximal spreading area for all DNabs formats at concentration of 6nM. **h,** Maximal spreading area for all DNabs formats at concentration of 16 nM. Each point represents one experiment, and is the median value of 20-100 cells. **i**, Schematics of the effect of glycocalyx shedding by Neuraminidase. **j**, Engagement rate for all DNabs at 6 nM with neuraminidase. **k,** Maximal spreading area for all DNabs (except 30cis) at 6 nM with neuraminidase. Boxplots correspond to median ± interquartile range. All statistical differences between paired tests are indicated with #.

The effect of DNab length on the engagement rate suggests that the glycocalyx may act as a steric barrier impeding engagement. To test this hypothesis, we preincubated T-cells with neuraminidase (also called sialidase), an enzyme cleaving negatively charged sialic acids from terminal positions of glycan chains, leading to a reduction in glycocalyx thickness and density, as schematically represented in **Fig. 2i** and controlled by flow cytometry (**Extended data Fig 7**). Treated cells engaged much faster for all DNab formats except 30cis (compare **Fig 2d** and **2j, Extended Data Fig. 6**). The treatment augmented the maximal spreading area in all cases, abrogating differences between formats (**Fig. 2k**). Therefore cell engagement was limited by the glycocalyx barrier, which hinders bridging by the shortest DNabs, and can be more easily crossed by longer constructs.

### Arrest of motile cells on ICAM-1 and antigen coated surface depends on DNab concentration and length

To better mimic the surface of a tumor cell, the adhesion molecule ICAM-1 was used together with HER2 to co-functionalize the surface (**Fig 3a**). Activated T lymphocytes are known to migrate on immobilized ICAM-1 by interaction via the integrin LFA-1^20^. However, in the presence of DNabs, they were forced to stop, leading to the establishment of a circular synapse (see representative Bright Field/RICM images in **Fig 3b**). The engagement rate, engaged fraction at 10 min, and maximal spreading area were determined as before. The first two parameters were found to be independent of DNab format and concentration (**Fig. 3c, Extended Data Fig. 8**), strongly suggesting that the initial engagement on the cell onto the surface was mediated by ICAM-1, in an engager independent manner. Consistent with this hypothesis, the spreading area at 6 nM DNab was found to be independent of DNab format, while nevertheless being slightly larger than on ICAM-1 alone (**Fig. 3d**). Once engaged, cells readily start to migrate until it may stop in the presence of DNab, establishing a circular synapse (**Fig. 3b, Supplementary Video 2**). Using cell tracking, a cell arrest was defined when the velocity went below a threshold of 3 µm/min. The distribution of the dwell times between the engagement onset and the immobilization was represented as a survival curve (see representative example of stop survival curve on with DNabs 30 or no DNab on **Fig. 3e**). The stop rate, defined as the slope of the stop survival curve, was measured as a function of DNab format, for several concentrations. In the absence of DNab or at low concentration (0.6 nM), cells practically did not stop migrating within the duration of the experiment (20-30 min). On the contrary, in the presence of 6 nM of any of the DNabs, cells rapidly stopped (**Fig. 3f, Supplementary Video 2**). At the intermediate DNab concentration of 1 nM, the format impacted the stop rate, being 10-fold slower for Dnab 40 than for DNab 20 (**Fig. 3g, Supplementary Video 2**). This trend was confirmed when considering the stopped fraction after 10 min of cell-surface engagement (**Fig. 3h**). Overall, the concentration of DNab strongly impacted the stop rate (by a factor 100 between 0.6-6 nM), and the DNab length had an impact within a narrow window around 1 nM.

**Fig. 3:**
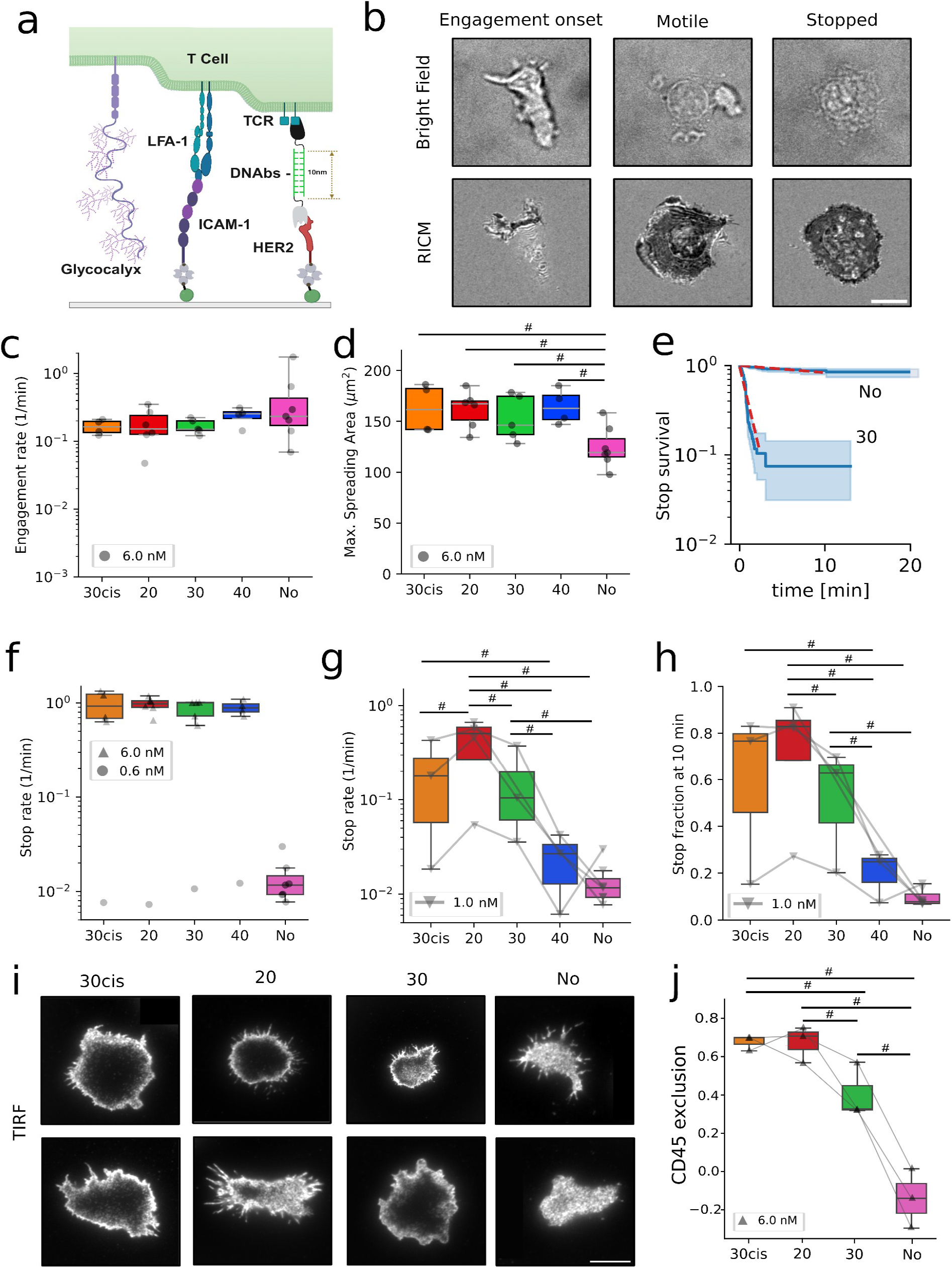
Cell-arrest on antigen surface with ICAM-1 is controlled by DNab concentration and format. **a**, Schematics of T-cell interaction with an antigen bearing surface with ICAM-1 in the presence of DNabs. **b,** Bright field and RICM micrographs of a cell at engagement onset, in motile or stopped phase. Scale bar: 8 µm. **c**, Engagement rate for all DNab formats at 6 nM concentration, or No DNab as a negative control. Each point represents one experiment with 20-100 cells. **d**, Maximal spreading area for 20, 30, 40 formats and No DNab as a negative control. Each point representing one experi ment, is the median of 20-100 cells. **e,** Representative curves (each representing 20-100 cells) of the stop survival before arrest and after cell engagement on the surface for No DNab and DNabs 30 at 6 nM. The slope of the dashed line defines the stop rate. **f,** Stop rate calculated from engagement time, as a function of the DNab format at concentrations of 0.6 and 6 nM, or control with No DNab. **g,** Stop rate calculated from engagement time, as a function of the DNab format at concentration of 1 nM, or control with No DNab. **h**, Fraction of stopped cells after 10 min, as a function of the DNab format at concentration of 1 nM. **i**, TIRF micrographs of cells on ICAM-1 alone, or with 6 nM DNab 30cis, 20, 30, fixed and stained for CD45. Scale bar: 8 µm. **j,** CD45 exclusion as a function of DNab format or on ICAM-1 alone (1 point per cell; pool of 3 independent experiments). Boxplots correspond to median ± interquartile range. All statistical differences between paired tests are indicated with #.

### CD45 exclusion depends DNab length

To explain why shorter DNabs were more efficient at arrest in the presence of ICAM-1, we conjectured that short engagers could lead more efficiently to the exclusion of the phosphatase CD45 from the synapse, leading to stronger cell activation^21^. To explore this hypothesis, CD45 was labelled after fixation of cells engaged on ICAM-1 and DNabs and the cell surface interface was imaged by TIRF. While CD45 is visible all under the cell in the absence of DNab, it was partially or largely excluded from the synapse in the presence of 6 nM of DNab (**Fig. 3i**). The phenomena was further quantified by measuring the fluorescence intensity in the middle of the cell and dividing it by the intensity at the edge. The exclusion was maximal for DNabs 30cis and 20, lower for 30 and absent in the absence of DNab (**Fig 3j**). Thus, in the presence of extra adhesion, short engagers can efficiently induce the exclusion of CD45, presumably leading to better effector function.

### Cytotoxic potency is determined by DNab length

To study the functional activity of DNabs, T cell-mediated cytotoxicity assays were conducted via live-cell imaging. All 4 DNabs (20, 30, 30cis, 40) promoted specific engagement between T cells and target MCF-7-HER2^+^ cells, inducing concentration-dependent lysis (**Fig. 4a, Supplementary Video** 3). Critically, no cytotoxicity was observed using DNabs nC and Hn, confirming that cell lysis required simultaneous engagement of both CD3 and HER2 (**Supplementary Table 2**).

**Fig. 4:**
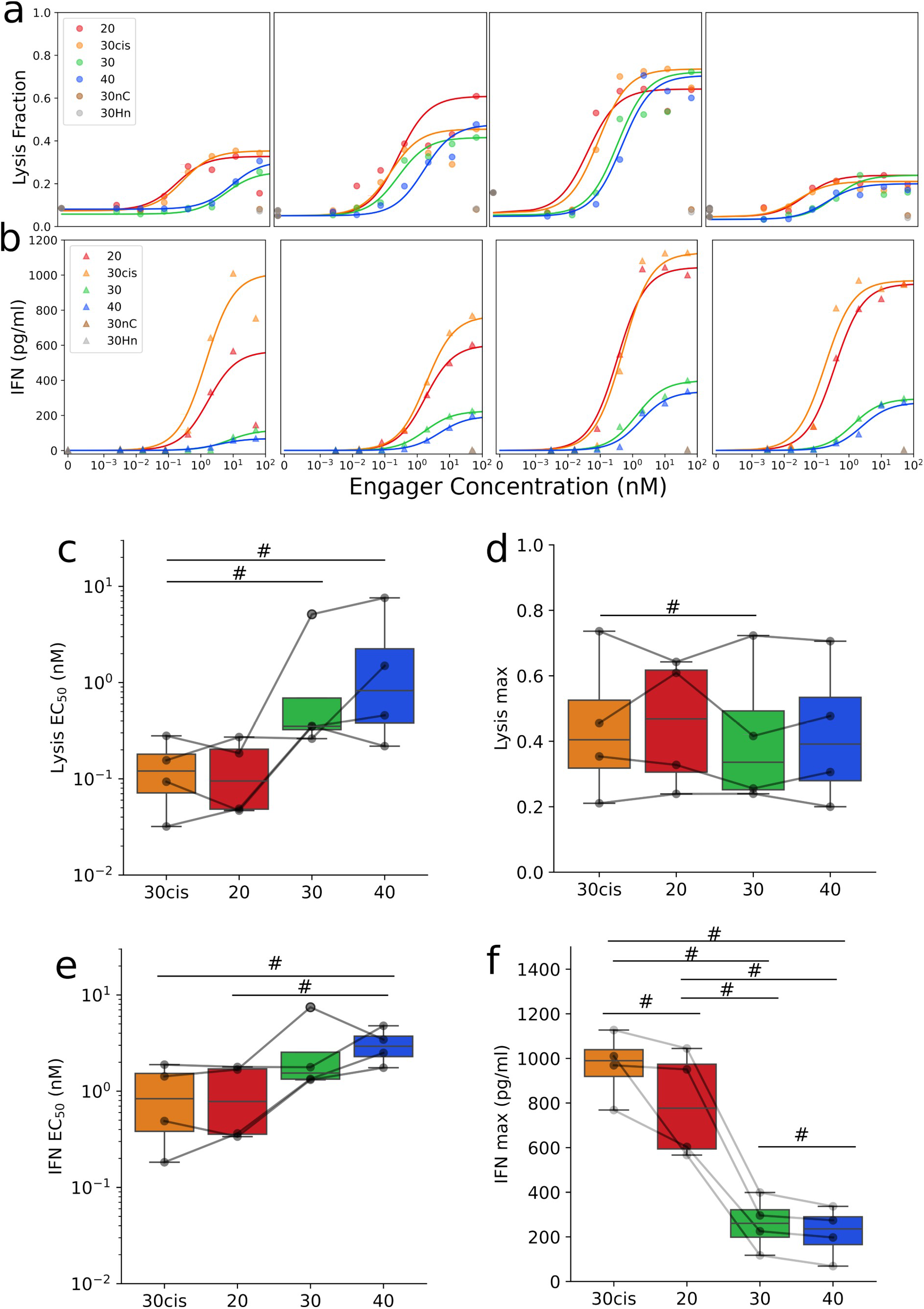
DNab-mediated cytotoxicity and T cell activation in MCF-7-HER2 co-cultures. MCF-7-HER2^+^ target cells were co-cultured with CFSE-labeled primary T cells from four independent donors (E:T ratio 5:1) in the presence of serial dilutions of DNabs (0.016–50 nM), n = 4. **a,** Dose-response curves displaying the fraction of target cell lysis after 4 h. Cytotoxicity was quantified via live-cell imaging by monitoring PI uptake in Hoechst-stained target cell nuclei, n =4. **b,** Dose-response assessment of IFN-γ release detected in the culture supernatant at the experimental endpoint (≈ 4 h). **c**, Measured EC_50_ for tumor cell lysis. **d**, Maximal fraction for tumor cell lysis. **e**, Measured EC_50_ for IFN-γ secretion. **f**, Maximal IFN-γ secretion. n = 4. Boxplots correspond to median ± interquartile range. All statistical differences between paired tests are indicated with #. *EC_50_: half-maximal effective concentrations; IFN-γ: interferon gamma*

We noticed an inverse relationship between construct length and cytotoxicity. DNab 20 exhibited the highest efficiency, achieving EC_50_ of 0.1 ± 0.1 nM, followed by the 30cis at 0.1 ± 0.1 nM. In contrast, the longer constructs showed reduced potency, with median EC_50_ values increasing to 1.5 ± 2.4 nM for DNab 30 and 2.4 ± 3.5 nM for DNab 40. A significant difference was observed between DNab 20 and 40, and between DNab 30cis and 40 (paired estimation test with 95% confidence) ( **Fig. 4c**)). Strikingly, DNab 20 was approximately 24-fold more potent than DNab 40, demonstrating that DNab length alone was sufficient to modulate the cytotoxic efficiency, and highlighting the critical role of synaptic tightness in T-cell activation.

### IFN-γ release is functionally decoupled from cytotoxic efficiency

We next compare the ability of DNabs to trigger cytokine secretion. While the EC_50_ regarding IFN-γ secretion was comparable across constructs, a drastic difference was observed in the amplitude of the response (**Fig. 4b**). The shorter DNabs (20 and 30cis) triggered a robust IFN-γ secretion, reaching maximal levels (Emax) of 791 ± 241 pg/mL and 969 ± 149 pg/mL, respectively. In striking contrast, increasing the length of the DNA linker led to a minimal IFN-γ release, plateauing at 259 ± 119 pg/ mL and 219 ± 115 pg/mL for DNabs 30 and DNabs 40 (**Fig. 4f**). Interestingly, these longer constructs maintained a robust lytic activity, reaching saturation levels similar to the 20 and 30cis formats, while resulting in a four-fold reduction in cytokine secretion levels compared to the short formats. All together these results suggest that cytotoxicity and cytokine release can be partially dissociated using only geometrical parameters.

### Trispecific DNabs as molecular OR-gates for multispecific engagement

Having evidenced some of the trends ruling bispecific DNab efficacy, we next decided to exploit the versatility afforded by DNA based linkers to generate trispecific T cell engagers targeting two different tumor antigens with the aim of alleviating one of the classical mechanisms of resistance displayed by tumor cells, i.e., tumor antigen loss. As a proof of concept, we decided to target HER2 and EGFR, two receptors often co-expressed by tumors. Trispecific constructs targeting CD3, HER2, and EGFR were designed to implement a Boolean ‘OR’ logic gate, enabling T-cell activation against tumor cells expressing either HER2 or EGFR, or both (**Fig. 5a**). These constructs were generated using 30 bases oligonucleotides and purified following the same protocol as bispecific DNabs. Their purity was verified by agarose gel electrophoresis (**Fig. 5b**).

**Fig. 5:**
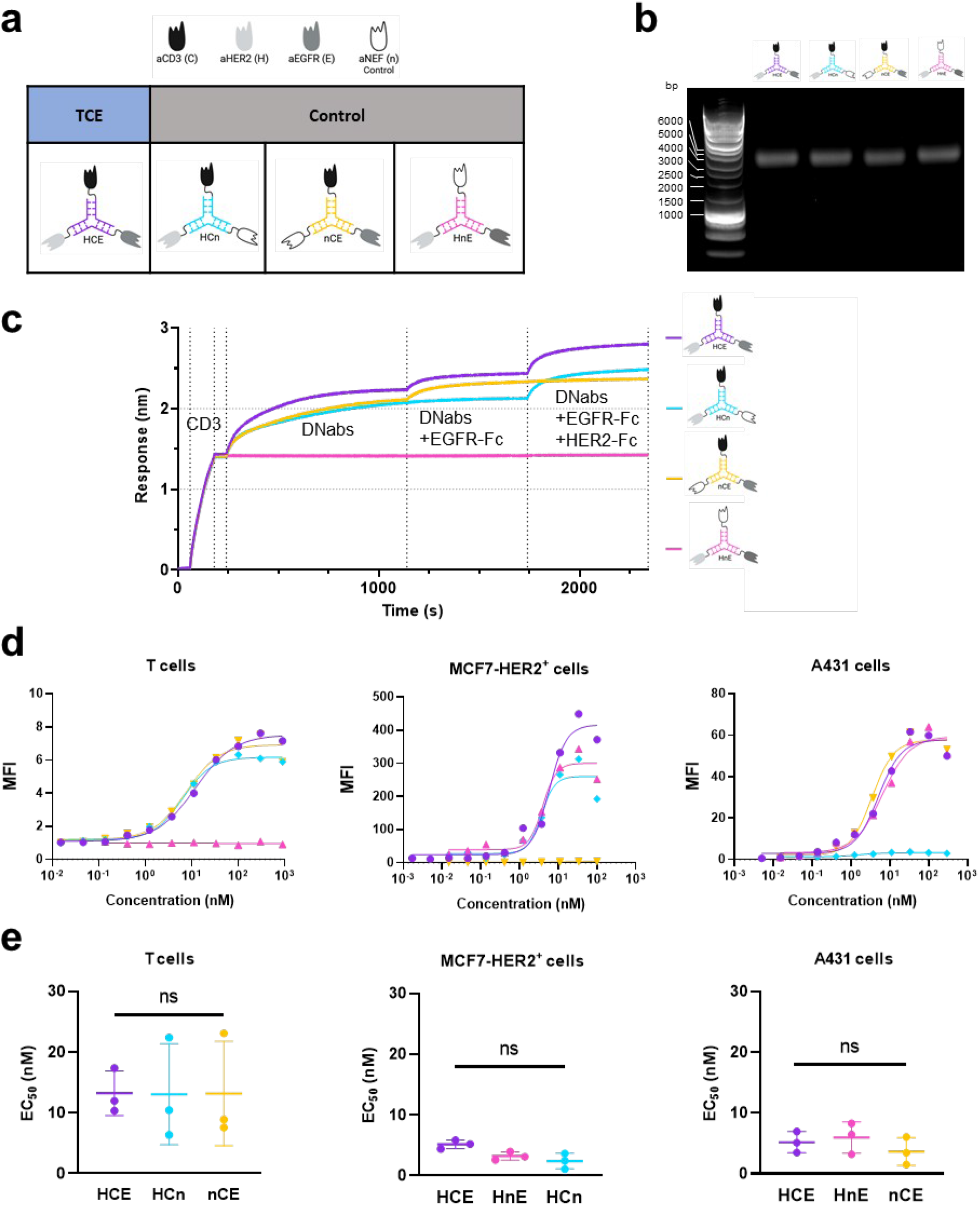
Design and characterization of trispecific anti-HER2 x CD3 x EGFR DNabs. **a,** Schematic representation of trispecific DNabs (HCE) and control constructs carrying anti-Nef19 (n) in place of one targeting domain (HCn, HnE, nCE). **b,** Agarose gel electrophoresis confirming construct identity and purity. **c,** Bio-Layer Interferometry (BLI) multi-step binding assay to validate the simultaneous binding of DNabs to their three targets, *n* = 3. Streptavidin biosensors pre-loaded with recombinant CD3 epsilon-delta were sequentially exposed to DNabs (50 nM), EGFR-Fc (10 μg/mL) +DNabs (50 nM), and finally HER2-Fc (20 μg/mL) + EGFR-Fc (10 μg/mL) + DNabs (50 nM). Stepwise binding profiles confirmed ternary complex formation. **d,** Representative binding curve obtained by flow cytometry on T cells, MCF-7-HER2^+^, A431 cells, showing apparent affinity (EC_50_), n = 3. **e,** Summary of EC_50_ values from flow cytometry experiments on three cell lines (T cells, MCF-7-HER2^+^, A431), n = 3 biological replicates per cell type; data presented as mean ± SD (respectively p = 0.7, 0.12, 0.09 One-Way ANOVA). *Abbreviations: aHER2: nanobody anti-HER2; aCD3: nanobody anti-CD3; MFI: Median Fluorescence Intensity*

The ability of the trispecific DNabs to simultaneously interact with CD3, HER2, and EGFR was demonstrated using BLI. As expected, control molecules displayed restricted binding profiles consistent with their design (**Fig. 5c, Extended Data Fig. 9**).

To assess whether the trispecific DNabs could recognize endogenous antigens in a physiological setting, flow cytometry titrations were performed on cell lines express ing the specific targets using MCF-7-HER2^+^ cells (HER2^+^), A431 cells (EGFR^+^) and primary T cells (CD3^+^) (**Fig. 5d**). Analysis of the binding revealed similar apparent affinities across all formats (**Fig. 5e**). Statistical assessment confirmed the absence of significant differences in binding strength between the constructs. These data indicate that the specific spatial arrangement of trispecific DNabs did not induce steric hindrance, preserving efficient recognition of all 3 epitopes in the cellular environment.

We first profiled the cell surface density of HER2 and EGFR across three cancer cell lines using quantitative flow cytometry, confirming a very low expression of EGFR and low expression of HER2 for MCF-7, a stronger expression of HER2 for the same cell line transfected with HER2, and a strong expression of EGFR with very low expression of HER2 for cell line A431 (**Supplementary Table 3. Table S4**). We subsequently evaluated the cytotoxic potency of the trispecific DNabs using real-time cell analysis (RTCA) over 48 h. As expected, the trispecific DNab targeting HER2, EGFR and CD3 induced a complete lysis across all three cell lines, demonstrating robust lymphocyte T activity independent of the individual antigen expression profiles, while the control DNab devoid of CD3 specificity did not generate specific cytolysis. ( **Fig. 6a**)

**Fig. 6.**
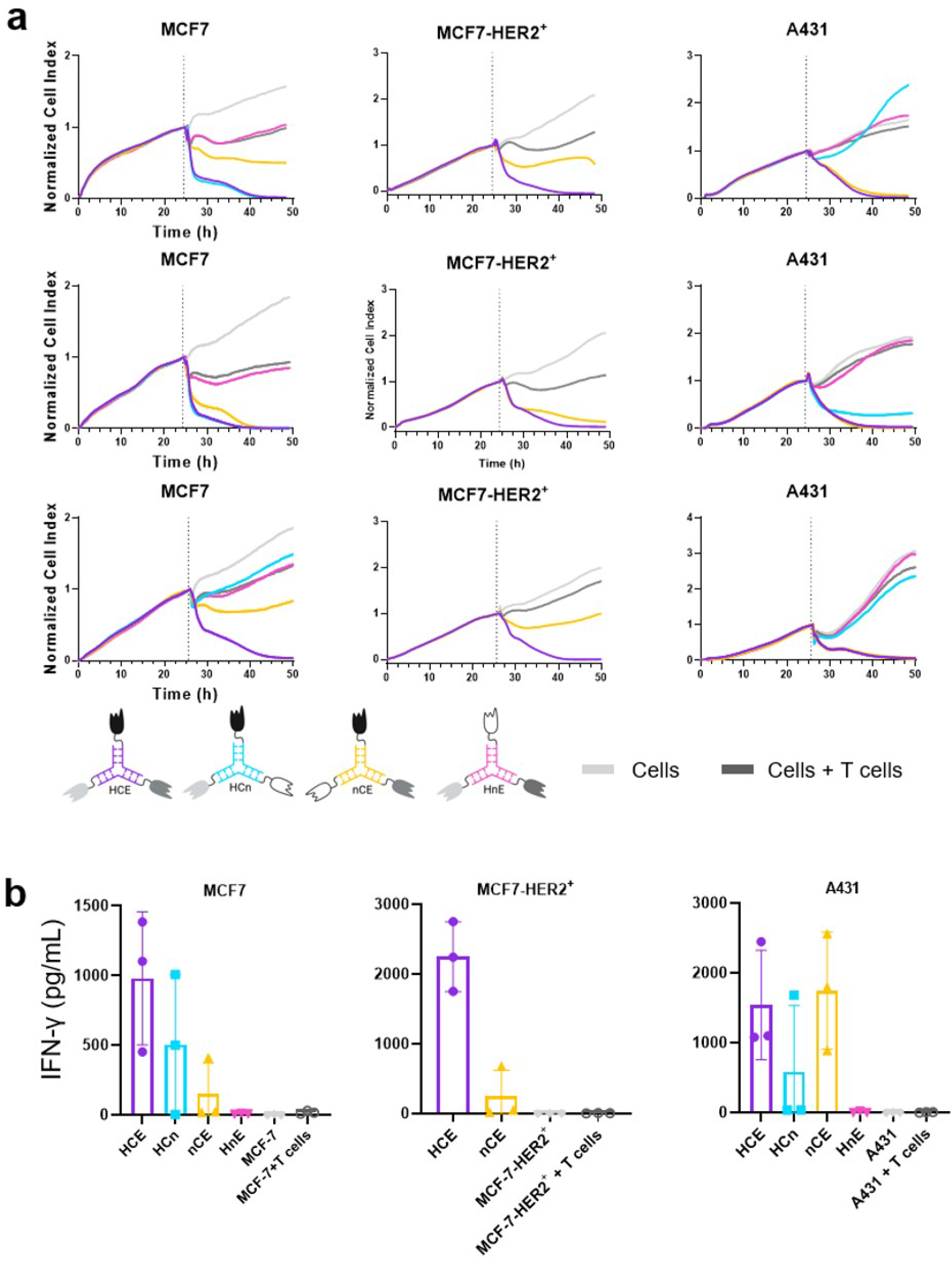
T cell-mediated cytotoxicity and cytokine secretion profile of trispecific DNabs. **a,** Realtime kinetics of tumor cell cytotoxicity monitored by impedance-based Real-Time Cell Analysis (RTCA). Target cells (MCF-7-HER2^+^, MCF-7, and A431 cells; 10,000 cells/well) were pre-cultured for 24 h to establish baseline impedance. After 24h (indicated by the vertical dashed line), trispecific DNabs (HCE, HCn, nCE, HnE; 10 nM) were simultaneously added with activated T lymphocytes [E:T; 5:1]). Cell Index was continuously recorded for 24 h. The stepwise decline in Cell Index reflects progressive target cell lysis. Representative cytotoxicity assay (*n* = 3) **b,** Quantification of interferon-gamma (IFN-γ) secretion. Cytokine concentration was measured by ELISA in culture supernatants collected at the end of the RTCA experiment (24h post-coculture). Data represent mean ± SD from three independent biological replicates using T cells from different donors (*n* = 3). *Abbreviations: IFN-γ: interferon-gamma*.

In contrast, the efficacy of control constructs lacking one of the specificities was governed by the expression of their respective antigens on target cells. While nCE targeting only EGFR achieved complete lysis (100%) on EGFR^high^ A431 cells, its potency was significantly reduced on cell lines with limiting EGFR expression, such as MCF-7-HER2^+^ and MCF-7, where it only induced a partial cytotoxicity (**Fig. 6a**). Similarly, HCn targeting only HER2 was fully active on MCF-7 cells (100% lysis) but induced low cytotoxicity on the HER2^low^ A431 cell line (**Fig. 6a**), confirming that efficacy was constrained by antigen availability. Quantification of secreted IFN-γ was performed to assess T cell activation and demonstrated a strong correlation with cytotoxic activity (**Fig. 6b**).

Collectively, these data establish a clear correlation between tumor antigen density and cytotoxicity, highlighting the capacity of this trispecific design to effectively target heterogeneous tumors via a Boolean ‘OR’ logic gate mechanism.

## Discussion

TCEs mediate their action through the formation of immune synapses, the quality of which condition the efficacy of target cell lysis and the extent of cytokine secretion. In this context, the structure of the engager, and especially the distance separating the two antigen-binding sites, is of prime importance since it directly impacts the intermembrane space. Conventional TCEs relying on full size antibodies bearing flexible hinge regions or fusion of antibody fragments using flexible peptidic linkers preclude a strict control of this parameter. In this work, we leveraged the inherent rigidity of double stranded DNA used as linker to generate nanobody-based T cell engagers of well-defined lengths, that we named DNanobodies or DNabs. We first established their ability not only to bind specifically to T cells and target cells but also to engage both cell types concomitantly. This co-engagement constitutes the crucial prerequisite for cytotoxic activity, leading to T cell activation and target cell lysis.

To further characterize the DNabs and dissect the mechanisms by which they mediate cell-cell engagement, we studied how they control the interaction of primary T cells with HER2 coated surfaces, using surface sensitive microscopy such as RICM^22,23^. After a variable period where the T-cells scan the surface^24^, spreading starts and reaches a maximal value within time-scale of minutes ^23^. When no additional adhesion ligands are present on the surface, the bridging is exclusively provided by the DNab, which acts both as adhesion molecule and activation signal. In this case, the dwell time before spreading onset as well as the maximal spreading area are primarily controlled by DNab concentration, and secondarily by DNab format, the longer ones providing earlier engagement higher spreading. We showed that the length dependence arises in these conditions because the T cell glycocalyx acts as a physical barrier that prevents short DNab bridging, while long DNab are less influenced by the presence of the repulsive barrier. Interestingly, this order of efficacy is reversed in the presence of ICAM-1 and the antigen on the solid surface. In this case, the large extracellular domain of the integrin LFA-1 expressed by T cells can bind to ICAM-1, resulting in instant adhesion and spreading for a majority of cells, in a DNab-independent way. However, T cells start migrate on the surface and DNabs only enter the game by forcing T cells to stop, which happens in a format- and concentration-dependent fashion. Shorter constructs better promote arrest, which is likely related to their capacity to activate the TCR through the establishment of more efficient synapses, probably through a more efficient CD45 exclusion from the synapse^25–27^. Our results show a strong impact of DNab concentration on stopping probability, with a narrow concentration range exhibiting format dependence, reminiscent of the behavior observed in membrane adhesion mediated by weak and soluble linkers^28^. It would be interesting to push these investigations to describe the cellular switch associated with the transition between the two concentration regime. In addition to the kinetic segregation effect, the impact of length may be more directly due to accessibility reasons, as discussed recently in the context of T cells^29^. Even in a purely passive system, the length of the linker in comparison to the membrane separation is a crucial parameter for membrane adhesion since the reactants residing on the two opposing membranes separated by a gap must touch each other for effective formation of bonds^30^. In fact, membrane separation and fluctuations impact both the on-rate and the off-rate of bond formation^30^. In cells, the membrane-separation is imposed by glycocalyx, and membrane fluctuation is provided by active filopodial “tiptoeing”^31^. When the adhesion is only DNab-mediated, longer DNabs are more easily accessible, thus enabling easy adhesion. Shedding the glycocalyx removes this advantage and abrogates length dependence of spreading metrics. In presence of ICAM-1 – itself relatively long, the initial spreading is likely dominated by ICAM-1/ LFA-1 bonds and becomes DNab length independent. The subsequent DNab-length-dependent stopping is likely to be an active process involving CD45 exclusion. Taken together, these results suggest a non-trivial linker-length dependence of T cell engagement and activation, even in the very early stages.

DNabs were next characterized in the cytotoxicity context. While all formats induce tumor cell lysis, we observed marked differences in potency. In a first approximation, the concentration behavior could be rationalized using an equilibrium model of bispecific antibodies bridging and induced cytotoxicity^32^, relating the potency with receptor surface density and affinities. This model introduced a molecular cooperativity parameter describing the ratio of bispecific binding in solution *vs* within the synapse binding, which could be related to DNab length, controlling the ability of the TCE to cross the glycocalyx barrier.

In this cellular context, an inverse correlation between cytotoxic potency and molecular size was observed. Similar to cell arrest, the efficiency of target cell lysis progressively decreased as size increased. Strikingly, a minimal variation in the inter-paratope distance, in the range of only a few nanometers (≈ 7 nm), resulted in an order-of-magnitude difference in the cytotoxic EC_50._

The sensitivity to architectural variations, both for cell-surface spreading assay and cell-cell cytotoxic assay, underscores the critical importance of spatial geometry and inter-membrane distance in orchestrating the immune response. The insights from the cell-surface assay in the absence of ICAM-1 could be relevant in a tumoral microenvironment with variable ICAM-1 densities^33^. While the importance of target cell glycocalyx in tumor resistance was previously emphasized^34^, we show here that effector glycocalyx may also be relevant. Our work contributes to rationalize approaches using sialidase-coupled T cell engagers^35,36^ to accelerate target recognition by the effector cell.

In cytotoxic assays, as well as cell-surface assays in the presence of ICAM-1, the shorter DNabs are found to be more efficient in terms of T cell arrest and CD45 segregation. This observation resonates directly with classical results showing that longer TCR-pMHC bonds are less activating^37^, also applicable for the interaction of CAR-T cells with their antigen^38^. In this line, the Kinetic Segregation Model^21^ postulates that T-cell activation is governed by the physical exclusion of bulky inhibitory phosphatases, such as CD45 (extracellular domain ≈ 40 nm), from the close contact zone of the synapse, favoring the phosphorylation of the TCR/CD3 complex. Our results suggest that the shorter bispecific constructs (20 and 30cis) mimic this natural spatial organization by imposing an inter-membrane distance short enough to efficiently exclude CD45, thereby favoring TCR phosphorylation. This is confirmed by the labelling of phosphatase CD45 which shows that exclusion from the synapse is maximal for 20 and 30 cis, and is reduced for DNab 30. These results are consistent with previous reports showing that the molecular size of a TCE significantly modulates the resulting lymphocyte response.^11,14^ More specifically, our findings align with the framework established by Leithner *et al.* corroborating that intermembrane distance serves as a primary regulator of effector response selectivity.^11^

Importantly, our findings highlight a crucial functional decoupling. We observed that longer DNabs (30 and 40) induce significantly reduced pro-inflammatory cytokine secretion compared to the shorter formats (20 and 30cis). This geometric compromise allows for a drastic four-fold reduction in IFN-γ release while maintaining high tumor cell lysis capacity. This partial decoupling suggests a way to maximize the therapeutic window of these engagers for clinical translation. Our results indicate that DNab 30 could offer a favorable balance, enabling an effective tumor cell elimination (sub-nanomolar cytotoxicity) while minimizing the risk of immune overactivation. We hypothesize that DNab 30 generates a brief activation reaching the threshold for cytotoxic activity, but fails to provide the sustained signaling required to drive the prolonged transcriptional programs necessary for IFN-γ production. This scenario aligns with findings from Faroudi *et al.*, who identified two distinct activation thresholds within the same CD8^+^ T cells: a lower threshold enabling rudimentary lytic synapse formation, sufficient for granule exocytosis, and a higher threshold requiring a mature synapse and sustained calcium signaling to induce IFN-γ production^39^. Our results suggest that a partial exclusion of CD45 strongly contributes to reducing IFN secretion, with only a limited impact on cytotoxicity.

While these findings establish a clear geometric rule in our setting, the next critical step is to evaluate how these defined geometries perform across a broader biological landscape. Importantly, it should be kept in mind that the inter-membrane space also strongly depends on the size of the tumor antigen, on the location of the targeted epitope (proximal or distal from the tumor membrane), or even by the angle used by the antibody fragment to bind this epitope, suggesting that the design of an optimal TCE may need to be adapted to each target. This is however completely realistic for DNab-based TCEs, which are fully modular in terms of all these design parameters.

Using the versatility of our DNA-based approach, we generated trispecific TCE targeting two tumor antigens often co-expressed by epithelial cancers, i.e. HER2 and EGFR. The characterization of our trispecific formats validated the implementation of a logic OR-gate strategy. We confirmed that trispecific DNabs were capable of engaging CD3, HER2, and EGFR either independently or simultaneously. This dual tumor-associated antigen recognition capacity translates into versatile cytotoxic activity, allowing the effective lysis of cells expressing exclusively HER2, as well as those expressing only EGFR, and leading to a very efficient lysis of double positive target cells. This characteristic is of major importance in the context of solid tumors often exhibiting strong intra-tumoral heterogeneity, where distinct subpopulations of cancer cells may express different antigens. By enabling the targeting of a broader population, trispecific TCE help overcome resistance mechanisms associated with antigen loss. Of note, because of the inherent modularity and self assembly of this trispecific format, it becomes straightforward to generate on demand various combinations of “off-the-shelf” nanobody-oligonucleotide conjugates targeting different antigens.

The modularity of our platform could also be leveraged to implement AND-gate logic. By incorporating binders with modest affinity, it would be possible to restrict binding to double positive tumor cells through an avidity effect, thereby sparing normal cells that weakly express either target. This approach would significantly reduce “on-target, off tumor” toxicities that often limit the clinical use of T cell engagers ^40^. Another promising future lies in the engineering of trispecific DNabs capable of delivering an additional co-stimulatory signal, such as CD28. This strategy, currently at the forefront of preclinical and clinical investigation, aims to spatially restrict co-stimulatory signals to T cells actively engaged by the TCE at the tumor site^41^.

## Acknowledgements

We thank Rémy Torro and Stéphanie Junoy for excellent technical support, as well as Pierre-Henri Puech for useful discussions. This work was supported by institutional funding from CNRS, INSERM, and Aix-Marseille University and by the grant “DNAnobodies” from the French National Agency for Research (ANR). The Molecular Motors and Machines team at IBENS is an “Equipe labellisée” by the Ligue Nationale Contre le Cancer.

## Material and Methods

### Cell lines and primary cells

The human cell lines A431 and MCF-7 were obtained from the American Type Culture Collection (ATCC). The MCF-7-HER2^+^ cell line, a variant of MCF-7 engineered to stably overexpress the HER2 receptor, was previously described by González Gutiérrez et al.^10^ A431 cells were maintained in DMEM, while SKBR-3, MCF-7, and MCF-7-HER2^+^ cells were cultured in RPMI-1640 medium. All media were supplemented with 10% fetal bovine serum (FBS). To ensure the stable maintenance of HER2 expression, the growth medium for MCF-7-HER2^+^ cells was further supplemented with 160 µg/mL Hygromycin B (Thermo Fisher Scientific, 10687010).

All cultures were maintained in a humidified incubator at 37 °C with 5% CO_2_. Cells were pass every 3–4 days upon reaching approximately 80 % confluency. The absence of mycoplasma contamination was confirmed on a monthly basis using the MycoAlert Mycoplasma Detection Kit (Lonza Bioscience, LT07-318).

Primary T cells were purified from peripheral blood mononuclear cells using the kit pan T cell isolation kit (Miltenyi Biotec,130-096-535). Following an activation period of 48 h in RPMI-1640+ 10% FBS medium supplemented T Cell TransAct at a ratio of 10 μL per 10^6^ cells (Miltenyi Biotec, 130-111-160), T cells were maintained in expansion media containing solely IL-2 at 50 ng/mL until the day of the experiment.

### Agarose Gel Electrophoresis

Analysis of DNabs structures was performed by electrophoresis on 1 % (w/v) agarose gels. Gels were prepared using UltraPure™ Agarose (Invitrogen, 16500-500) in Tris-Acetate-EDTA (TAE) buffer (Merck, LSKMTAE50). Samples were mixed with TriTrack DNA Loading Dye (Thermo Fisher Scientific, R1161) prior to loading. A Smart Ladder (Eurogentec, MW-1700-10) was used as a molecular weight marker. Electrophoresis was conducted at 100 V for 60 min in TAE buffer. DNA bands were visualized using SYBR™ Gold Nucleic Acid Gel Stain (Life Technologies, S11494).

### Production and purification of the nanobodies

Nanobodies targeting Her2 (HER2.CE4), EGFR (D10), and the irrelevant Nb targeting Nef19 were generated in previous studies.^42–45^ The anti-CD3 V_H_H sequence was sourced from patent US20190382485A1. The phagemid of specific clones were transformed into BL21DE3 E. Coli. The production was induced by culturing the bacteria with Isopropyl β-D-1-thiogalactopyranoside 100 μM at 30 °C overnight. The culture was pelleted and lysed using Bugbuster lysis buffer (Merck Millipore) supplemented with benzonase (25 U/mL) and lysozyme (20 μg/mL). The his-tagged Nb-containing lysate was loaded on TALON^®^ SuperflowTM resin (Cytiva, 28957502). After two successive washes in PBS 300 mM NaCl followed by PBS, Nbs were eluted in PBS Imidazole 250 mM. Imidazole was removed using PD-10 desalting column (Cytiva, 17085101) and Nbs were stored in PBS. Protein concentrations were quantified via absorbance measurements at 280 nm using a NanoDrop One spectrophotometerprovider. To assess purity, samples were resolved under non-reducing conditions using 4–20 % Mini-PROTEAN TGX Stain-Free precast gels (Bio-Rad, Cat# 4568084). The Unstained Precision Plus Protei lnadder (Bio-Rad, 1610363) was used as the molecular weight marker for electrophoresis.

### Preparation of the nanobody-oligonucleotide conjugates

Nbs were site-specifically labeled using a chemo-enzymatic approach. First, bacterial transglutaminase (Zedira, T300) was used to catalyze the formation of a covalent bond between the glutamine of C-terminal c-myc tag of Nbs and the amine group of the reagent NH2-PEG3-Azide3 (Vector Laboratories: AZ101-1000). Subsequently, the azide-functionalized nanobodies were purified by gel filtration Superdex 75 10/300 GL column using an ÄKTA system (Cytiva) and conjugated to dibenzocyclooctyne DBCO-modified oligonucleotides via copper-free strain-promoted azide-alkyne cycloaddition (SPAAC) click chemistry.

### Production of the DNanobodies

Nb-oligonucleotide conjugates carrying complementary strands were mixed at an equimolar ratio and incubated at room temperature for 3 h to allow hybridization. The resulting complexes were purified by gel filtration on a Superdex 200 10/300 GL column (Cytiva) equilibrated in PBS. Eluted fractions were analyzed by both agarose gel electrophoresis and SDS-PAGE. Pure fractions were pooled, and the final concentration was determined using a NanoDrop spectrophotometer based on DNA absorbance. The molar concentration was calculated using the combined molecular weight of the hybridized oligonucleotide duplex.

Oligonucleotide sequence:

20: Unp1_20: 5’DBCON-AACCTTTTCCCTGCGTATCG-3’ and Unp2_20: 5’DBCON-CGATACGCAGGGAAAAGGTT-3’

30cis: Unp1_30: 5’DBCOTEG-ATATGAGGCGGTTATTCTTCGGCTCTTACA-3’ and Unp2_30cis: 5’-TGTAAGAGCCGAAGAATAACCGCCTCATAT-DBCON3’

30: Unp1_30: 5’DBCOTEG-ATATGAGGCGGTTATTCTTCGGCTCTTACA-3’ and Unp2_30: 5’DBCOTEG-TGTAAGAGCCGAAGAATAACCGCCTCATAT-3’

40: Unp1_40: 5’DBCON-GAGAATGGCCTCGCGGAGGCATGCGCCATGCTAGCGTGCG-3’ and Unp2_40 5’DBCON-CGCACGCTAGCATGGCGCATGCCTCCGCGAGGCCATTCTC-3’

For trispecific constructs, three distinct Nb-oligonucleotide conjugates were mixed at an equimolar ratio to allow self-assembly. Hybridization, gel filtration, agarose gel and SDS-PAGE were performed as described above. The final concentration was determined spectrophotometrically based on DNA absorbance, using the combined molecular weight of the three oligonucleotide strands (Unp2_30: 5’DBCOTEG-TGTAAGAGCCGAAGAATAACCGCCTCATAT-3’ Unp3_30: 5’DBCON-ATATGAGGCGGTTATTGCAACCGATACATT-3’ Unp4_30: 5’DBCON-AATGTATCGGTTGCATCTTCGGCTCTTACA-3’)

### Bio-Layer Interferometry (BLI)

Binding kinetics were characterized using bio-layer interferometry on an Octet® R2 system (Sartorius). The following proteins were used as ligands: recombinant human commercially biotinylated HER2 (H82E2 Acrobiosystems) and CD3 epsilon-delta (CT038-H2508H Sino Biological). Assays were performed at 25 °C. Streptavidin (SA) biosensors were pre-hydrated for 10 min in assay buffer (PBS, 1 % BSA, 0.05 % Tween 20). The assay consisted of four steps : i) Loading, ii) Equilibration, iii) Association, and iv) Dissociation. I) Loading: Biotinylated ligands were immobilized onto the sensors for 120 s. Optimization determined the following loading concentrations: HER2 (3 µg/mL), CD3 epsilon delta (1.25 µg/mL) ii) Equilibration: Sensors were equilibrated in buffer for 60 s to establish a stable baseline and remove loading excess. iii) Association: Sensors were dipped into wells containing graded concentrations of the DNabs (20, 30cis, 30, 40) for 300 s. The DNabs were tested at concentrations ranging from 1.8 to 150 nM on HER2 and 1.8 nM to 50 nM on CD3 in a three-fold dilution series. Control DNabs (Cn and nH) were evaluated exclusively at the highest concentration. iv) Dissociation: Sensors were moved back to the buffer for 300 s to monitor the dissociation rate. Raw data were processed using Octet Analysis Studio software (v12.2). Data analysis was performed using a standard 1:1 Langmuir interaction model with global fitting, yielding the association (ka_)_ and dissociation (kd_)_ rates adalongside the calculated equilibrium constant (K_D_ = koff/kon). However, for interactions where binding curves deviated from a standard 1:1 kinetic profile, affinity was determined using steady-state analysis: Req = Rmax×[DNabs]/(K_D_+[DNabs])

To verify the simultaneous binding of the DNabs to their targets, a sandwich assay setup was employed. Biotinylated recombinant CD3 epsilon-delta (1.25 µg/mL) was immobilized on streptavidin biosensors for 60 s, followed by a 60 s baseline step. The sensors were exposed to DNabs 20, 30cis, 30, 40 or control DNabs Cn and nH (50 nM) for 900 s to allow capture. Subsequently, the sensors were immersed in a solution containing recombinant HER2-Fc (20 µg/mL) with the same DNabs used in the previous step for 400 s to monitor the secondary binding event.

To assess the ability of the molecule to engage three targets simultaneously, a multi-step binding assay was performed. Biotinylated recombinant CD3 epsilon-delta (1.25 µg/mL) was loaded onto streptavidin sensors for 120 s, followed by a 60 s baseline stabilization. The sensors were then subjected to a sequential association protocol: i) incubation with DNabs HCE, HCn, nCE, HnE (50 nM) for 900 s, ii) exposure to EGFR-Fc (20 µg/mL)+ DNabs (50 nM) for 600 s, iii) exposure to HER2-Fc (20 µg/mL)+ EGFR (10 µg/mL) + DNabs (50 nM) for 600 s. Binding responses were monitored in real-time to confirm the cumulative assembly of the complex.

### Flow cytometry

Flow cytometry experiments were conducted using a MACSQuant® X cytometer (Miltenyi Biotec). All staining steps were performed in V-bottom 96-well plates using PBS containing 2 % BSA as staining buffer, with antibodies added at manufacturer-recommended concentrations. Cell surface binding of the unlabeled constructs was evaluated using an indirect detection method. Cells were incubated with DNabs for 1 h at 4 °C and washed, followed by incubation with a Mouse anti-His antibody (1:1,000; Novagen, 70796-3) for 1 h at 4 °C and wash, and Goat anti-Mouse IgG(H+L)-AlexaFluor 647 secondary antibody (1:300; Invitrogen, A21235) for 45 min at 4 °C and wash. Bispecific constructs binding was assessed on HER2-positive MCF-7-HER2^+^ cells and primary human T cells (bispecific DNabs) and EGFR-positive A431 cells (trispecific DNabs). Target expression on tumor cell lines was validated by staining with anti-HER2 or anti-EGFR reference antibodies. HER2 expression on MCF-7-HER2^+^ cells was assessed with Brilliant Violet 421™ anti-human CD340 (HER2), clone 24D2 (BioLegend, 324420). For A431 cells, an anti-EGFR antibody Cetuximab (Bio X Cell, clone C225) was used. Detection was performed using a secondary Goat anti-Human IgG (H+L) antibody conjugated to Alexa Fluor 647 (Invitrogen, A21445). Primary human T cells were phenotyped using a multicolor panel of directly conjugated monoclonal antibodies (all from Miltenyi Biotec): anti-CD3 (VioGreen™, clone REA613), anti-CD4 (FITC, clone REA623), anti-CD8 (VioBlue®, clone REA734), anti-CD25 (PE-Vio®770, clone REA945), and anti-CD69 (APC, clone REA824). All antibodies were incubated for 45 min at 4 °C in the dark. Data Analysis : Single-cell populations were gated based on Forward Scatter (FSC) and Side Scatter (SSC) to exclude debris and doublets. Data analysis was performed using Flowlogic v7.3.

### Quantification of cell surface receptors density

Surface expression of HER2 and EGFR was assessed by flow cytometry on MCF-7, MCF-7-HER2^+^, and A431 cell lines. For HER2 staining, 2×10^5^ cells were incubated with the mouse anti-HER2 antibody (clone 9G6, ref sc-08, Santa Cruz Biotechnologies) at a concentration of 10 µg/mL for 10 min at room temperature. Cells were then washed and incubated with a polyclonal antibody anti-mouse IgG FITC coupled (reagent 3 from Mouse IgG calibrator kit ref CP051 Biocytex). For EGFR staining, cells were incubated with the anti-EGFR antibody Cetuximab (Bio X Cell, clone C225) at 10 µg/mL, followed by a polyclonal antibody anti human IgG-FITC (reagent 3 from Human IgG calibrator kit, Biocytex CP010). To determine the absolute number of receptors per cell (Specific Antibody Binding Capacity, SABC), calibrations kits, Mouse IgG calibrator kit (Biocytex CP051) and Human IgG calibrator kit (Biocytex CP010), were used according to the manufacturer’s instructions. Calibration beads were stained in parallel with the cells using the same secondary antibodies and acquisition settings.

Data were acquired on a cytometer Fortessa X20 (BD Biosciences) and analyzed using FlowJo (BD Biosciences). The expression level of the tested antigen is determined using a calibration curve and reported as Specific Antibody Binding Capacity (sABC).

### Primary T-Cell glycocalyx removal

For some experiments, the glycocalyx of T cells was treated as follows. One mL of medium containing around 1 million cells was removed from the culture flask for experimentation. These cells were centrifuged and resuspended in 500 µl of serum-free RPMI medium. Neuraminidase (Purified, Worthington Biochemical, LS004761) was added to a final concentration of 1 unit/mL, and cells were incubated for 1 h at 37 °C. Following incubation, 500 µL of complete RPMI medium with serum was added to block neuraminidase activity, cells were centrifuged again, and resuspended in 1 ml of complete medium. 100 µl of cell were used for flow cytometry by adding 400µl of complete medium and 1µl of 5 mg/ml WGA-Rhodamine (Vector Laboratories, RL-1022-5). Samples were incubated for 5 min, away from light, and analysed on a cytometer Fortessa X20 (BD Biosciences).

### Cell spreading assay

For all cell spreading experiments, uncoated eight-well chambered polymer coverslip (Ibidi, 80801) were first incubated with 200 µl of 100 µg/mL BSA-biotin (Sigma-Aldrich, A8549) in PBS for 30 min at room temperature (RT) under agitation, followed by five times PBS rinses. Next, 10 µg/mL NeutrAvidin (Thermo Scientific, 31000) in PBS was added for 30 min at RT with agitation, followed by another 5 PBS rinses. Surface were then functionalized with 15 nM solution of biotinylated HER2 protein (Sino Biological, 10004-H27H-B) or when indicated, with a mixture of 15 nM of biotinylated HER2 and 90 nM of biotinylated ICAM-1 (Sino Biological, 10346-H49H-B) followed by PBS rinsing, Sample were transferred under the microscope chamber at 37 °C. Prior to Effector primary T cells (∼3 × 10⁴) addition in the chamber the DNabs were introduced at the desired concentration taking into account the final volume after addition of cells.

Cell spreading dynamics was monitored with time-lapse Reflection Interference Contrast Microscopy (RICM) using an inverted microscope equipped with 63x antiflex Zeiss objective (NA = 1.25), a green LED light source (λ = 546 nm) and a 14-bit CCD detector (Andor iXonEM, Oxford Instruments). For each condition multiple fields of view (9 to 16) were recorded through cyclic imaging over a 15-20 mins with temporal resolution of 15-20 s.

Time-lapse sequences were analyzed using the open-source software Celldetective^19^, which entails image preprocessing, segmentation, cell tracking, and time dependent signal extraction (e.g., spreading area, RICM intensity, velocity). From these analyses, key temporal parameters were extracted such as time of first contact detection, the onset of spreading, the arrest of motile cells by setting threshold on the extracted features. Details are provided in Supplementary information.

### Immunofluorescence after spreading / TIRF microscopy

For immunofluorescence labeling, cells were allowed to spread on functionalized surfaces as described above for 15 min. Fixation was performed with 2 % paraformaldehyde (PFA, Sigma) in PBS for 10 min at 37°C, followed by gentle PBS washes, taking care to minimize cell detachment throughout the procedure. Cells were incubated for 20-30 min in blocking buffer (1 % BSA in PBS) . Cells were incubated with anti-CD45 antibody (clone REA747, Miltenyi Biotec, Vio R667) diluted 1:200 in blocking buffer for 30 min at room temperature. Samples were rinsed four times with PBS and proceed with imaging.

Total internal reflection microscopy (TIRFM) and reflection interference contrast microscopy (RICM) were performed using an inverted microscope (Axio Observer, Zeiss), equipped with an EMCCD camera (iXon, Andor ). TIRF and RICM images were taken with a custom 100 x 1.46 NA oil antiflex Zeiss objective. For each condition, more than 30 cells were imaged across multiple fields of view to ensure reproducibility and statistical relevance.

Analysis was performed using Celldetective (ref 19), including background subtraction, and segmentation. A normalized distance transform approach was used to compute different zones within the cell mask: inner, middle and outer zones. Mean fluorescence intensity was measured for each region. CD45 exclusion was quantified using the ratio of inner to outer intensity and expressed as 1(I_inner/ I_outer) to reflect relative depletion from the center.

### Cytotoxicity under microscope

This protocol was adapted from the method described by Diaz-Bello et al.^18^; Target cells (MCF-7-HER2^+^) were seeded at a density of 20,000 cells per well in 96-multiwell plates (Nunclon Delta Surface, #167008) and allowed to adhere and proliferate overnight (18 h) under standard culture conditions (37 °C, 5 % CO₂, 85 %–95 % humidity). Prior to the assay, target cell nuclei were stained with Hoechst 33342 (5 µg/ mL) for 15 min at 37 °C. Cells were then rinsed three times with warm supplemented RPMI medium to remove excess dye. Effector primary T cells were labeled with CellTrace™ CFSE to allow for distinction from target cells. Briefly, T cells were incubated with CFSE (0.2 µL/mL per 1×10^6^ cells) in DPBS for 20 min at 37 °C. The reaction was quenched by adding 5 volumes of serum supplemented RPMI media, followed by a 5-min incubation at 37 °C. Cells were pelleted (600 g, 5 min) and resuspended in assay medium. The assay was initiated by removing the supernatant from target cells and adding Propidium Iodide (PI) at a final concentration of 4 µg/mL. Unlabeled DNabs were added to the culture at final concentrations ranging from 50 nM to 0.016 nM. Finally, the pre-activated T cells were added at an Effector:Target ratio of 5:1

Time-lapse imaging was performed using a Zeiss Axio Observer.Z1 inverted microscope equipped with a thermostated chamber maintained at 37 °C and 5 % CO₂. Images stacks were acquired every 15 min for a total duration of 4 h using a Plan-Apochromat 20× (NA 0.8) objective. Four channels were captured: Brightfield, Hoechst (Target nuclei), CFSE (Effector cells), and PI (Cell death marker).Quantitative image analysis was performed using using the open source Celldetective software. The workflow consisted of the following steps: (i) Segmentation: Target cell nuclei were automatically segmented in the Hoechst channel using a dedicated deep learning model (MCF-7_nuc_stardist_transfer) trained to accurately select target cell nuclei while excluding smaller effector cell nuclei. (ii) Feature Extraction: For each tracked nucleus, morphological and intensity features were measured at every time point, specifically the nuclear area and the mean fluorescence intensity. (iii) Event Detection: Cell death was identified utilizing the “Detect Event” module. A lysis event was defined as a sharp decrease in nuclear size (pyknosis) concomitant with an increase of propidium iodide and Hoechst decrease in Hoechst signal fluorescence intensity.

### Real-Time Cell Analysis (RTCA)

Cytotoxicity was monitored in real-time using the xCELLigence® RTCA system (Agilent). Target cells (MCF-7-HER2^+^, A431, and MCF-7) were seeded at a density of 10,000 cells/well in E-Plate 16 (Agilent, 300600870) and allowed to adhere and proliferate for 24 h. After the initial incubation, the assay was initiated by adding the pre-activated T cells at an Effector:Target (E:T) ratio of 5:1. Simultaneously, DNabs were added to the co-culture at a final concentration of 10 nM. Cell index (CI) values, reflecting cell viability and adhesion, were monitored continuously for 24 h.

### Cytokine Quantification by ELISA

Upon completion of the cytotoxicity assay (24 h after the addition of T cells and DNabs), cell culture supernatants were harvested to assess cytokine secretion. Interferon-gamma (IFN-γ) concentrations were determined using commercial ELISA kits (Thermo Fisher Scientific, 88-7316-88) according to the manufacturer’s instructions. Samples were diluted in PBS prior to analysis. Supernatants were diluted 5-fold (1:5), with a standard curve ranging from 3.9 to 500 pg/mL. Absorbance was measured at 450 nm using a microplate reader.

### Statistics

For figs. 2, 3, and 4, data were analyzed by the estimation statistics using the following package (Data Analysis using Bootstrap-Coupled ESTimation, https://github.com/ACCLAB/DABEST-python). Where ever possible paired test between two conditions (paired by day) with a confidence interval of 95% was performed, except if a condition would have less than 3 replicates. To determine half-maximal effective concentrations (EC50), dose-response curves were fitted to a Hill function of slope 1

Other data analysis was performed using the GraphPad Prism software. .

---

**Extended Data Figure 1.**
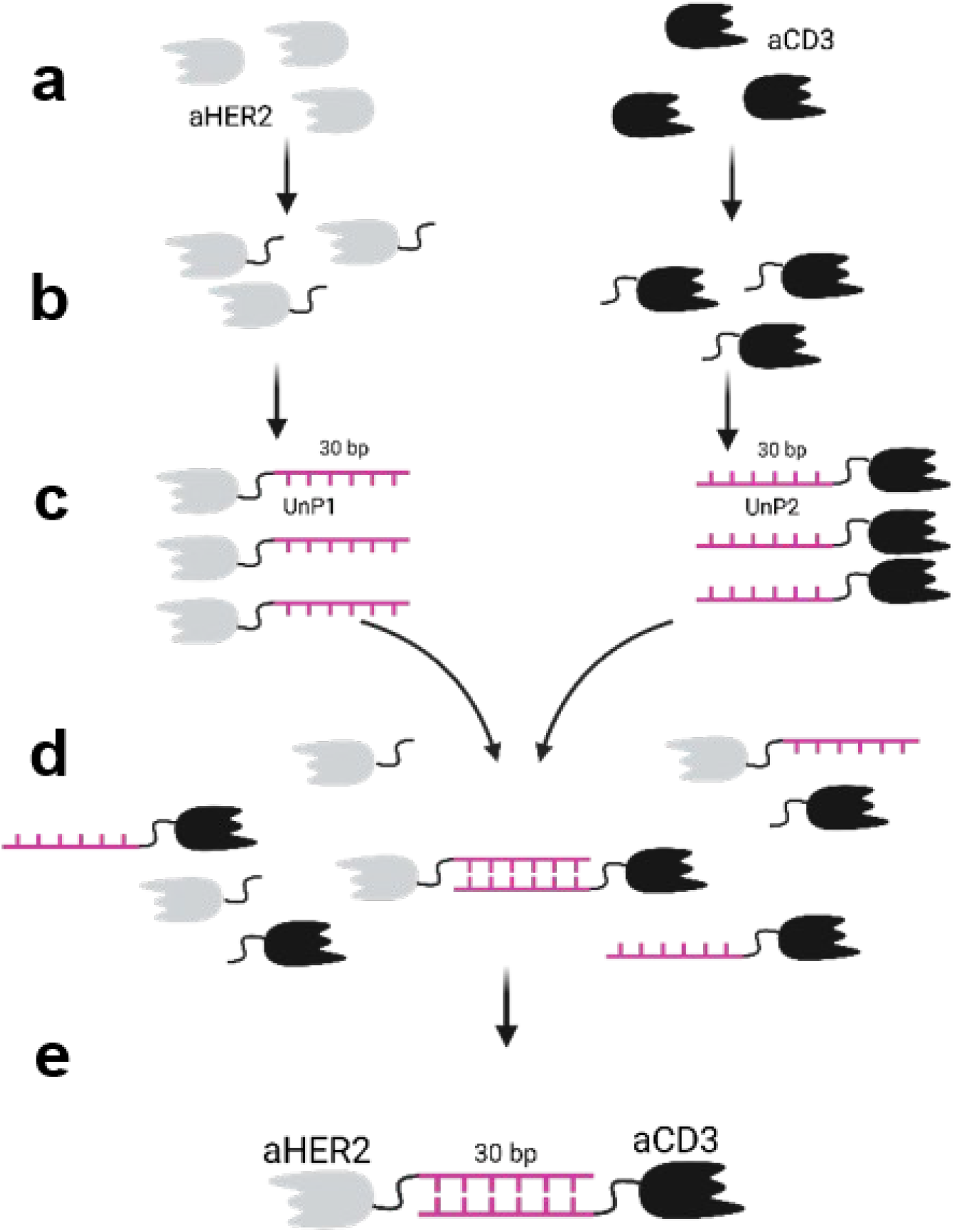
Schematic representation of the DNab assembly protocol. Example with DNAb 30. **(a)** Production of anti-HER2 and anti-CD3 nanobodies. **(b)** Site-specific conjugation via covalent bond formation between the C-terminal glutamine acyl group of the c-myc tag and the amine group of the NH₂-PEG₃-N₃ linker, followed by gel filtration purification to remove excess azide. **(c)** Click chemistry-mediated conjugation of UnP1 and UnP2 oligonucleotides to the nanobodies using a 1:2 molar ratio (oligonucleotides : nanobodies). **(d)** Equimolar hybridization of the two nanobody-oligo conjugates (HER2-UnP1 and CD3-UnP2). **(e)** Followed by a final purification step by gel filtration purification to remove excess components and isolate the pure DNabs.

**Extended Data Figure 2.**
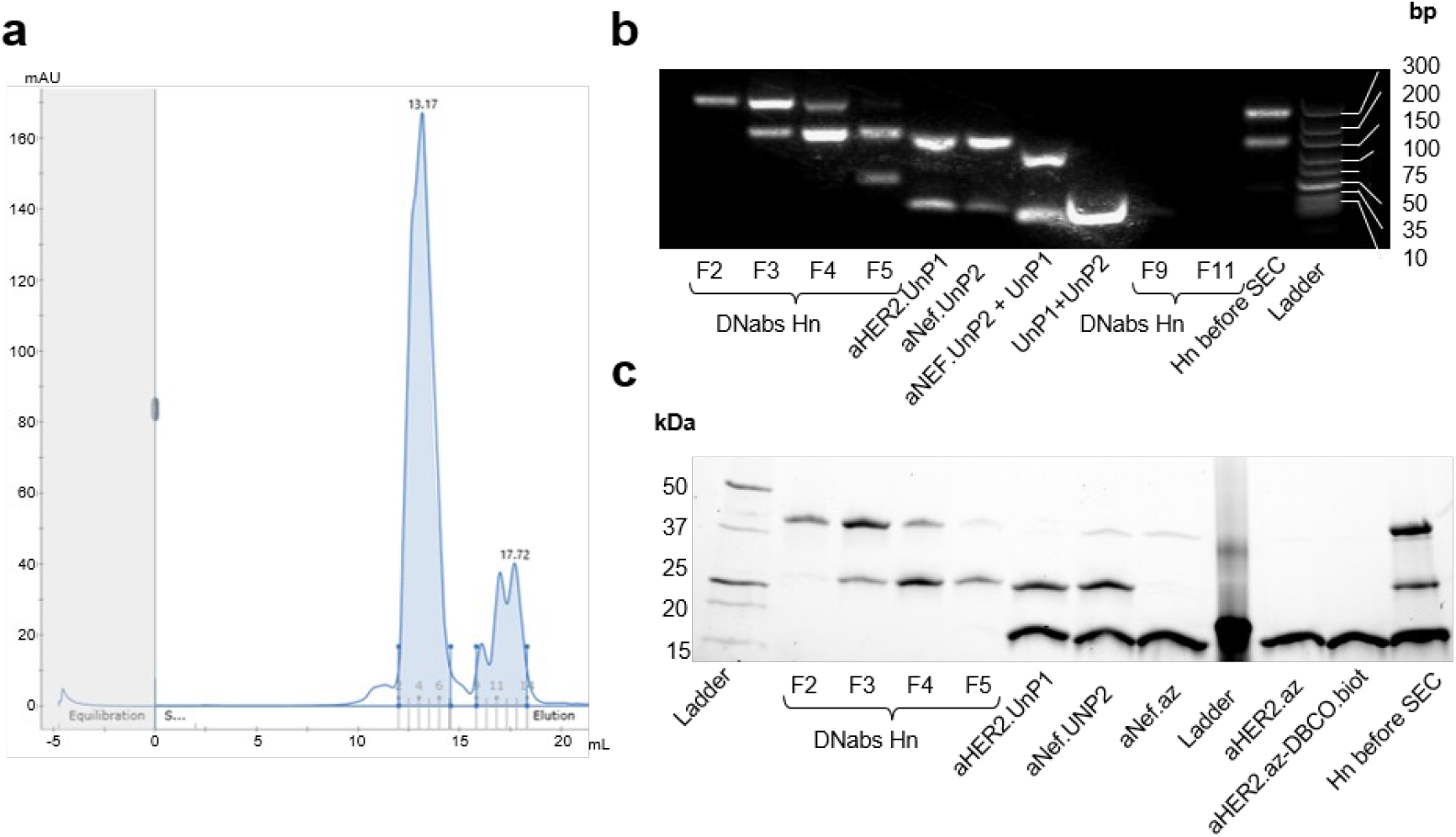
Purification of the anti HER2 x Nef DNab construct. **(a)** Gel filtration of DNabs on a Superdex 200 (Cytiva). Fractions corresponding to the major peak (fractions 1–4) and to the smaller secondary peak (fractions 9–11) were collected for further analysis. **(b)** Native agarose gel electrophoresis of purified DNab fractions and controls. **(c)** SDS PAGE analysis of the same samples as in **(b)**, with additional lanes containing monomeric HER2 nanobodies and Nef nanobodies (≈15 kDa).

**Extended Data Figure 3.**
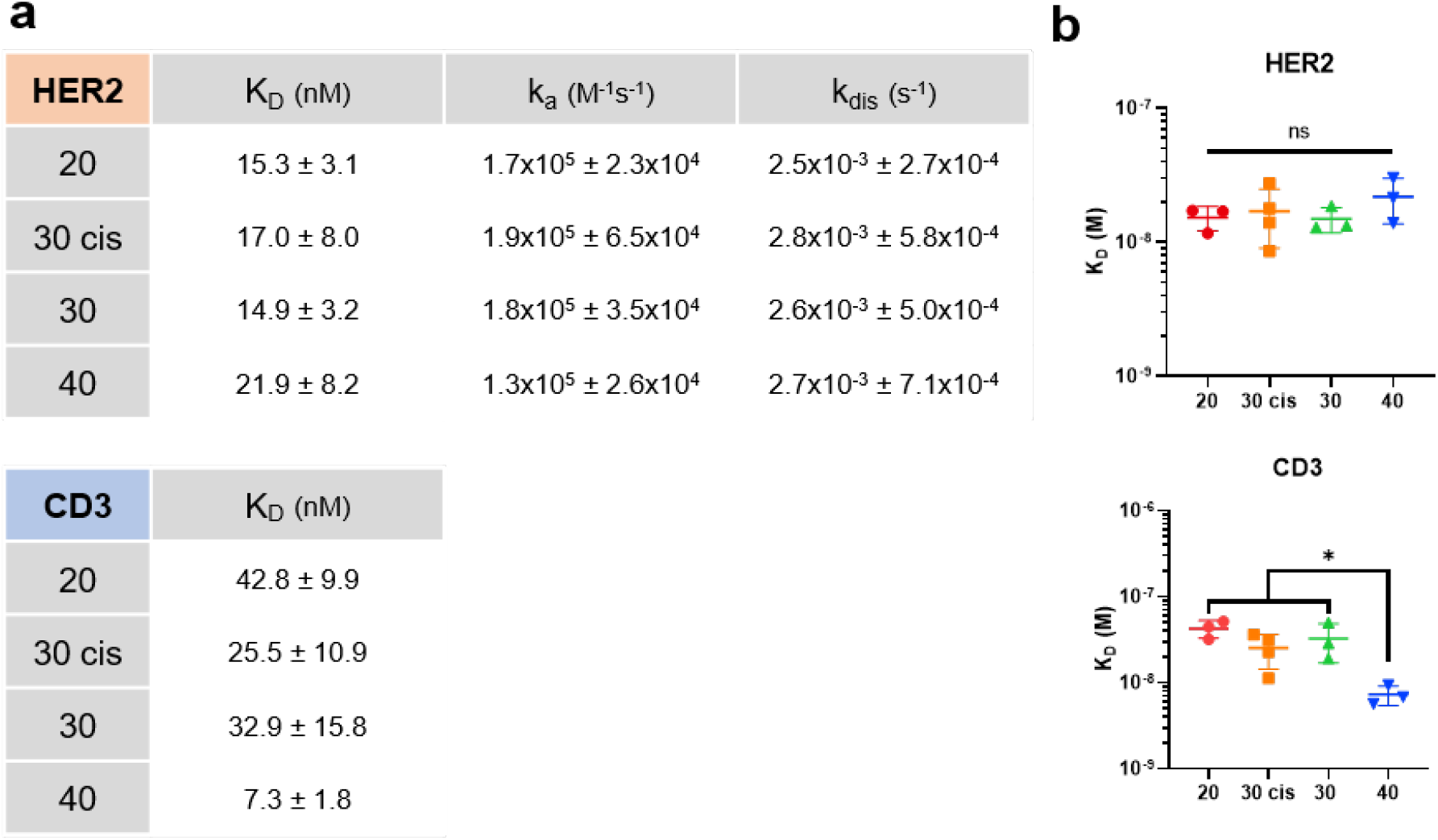
Binding affinities and kinetics of DNabs determined by bio-layer interferometry (BLI). **(a)** Summary table of kinetic and affinity constants measured using recombinant HER2 and biotinylated CD3εδ. For HER2, the association rate (kₐ), dissociation rate (kdis), and equilibrium dissociation constant (KD) were calculated using a 1:1 binding model. For CD3, KD was determined by steady-state analysis. Values represent the mean ± standard deviation (SD) from n ≥ 3 independent experiments. **(b)** Log scale plot of KD derived from the same BLI dataset as in **(a)**. Statistical analysis: One-way ANOVA followed by Tukey’s post-hoc test. ns = not significant (p > 0.05); * = p < 0.05.

**Extended Data Figure 4.**
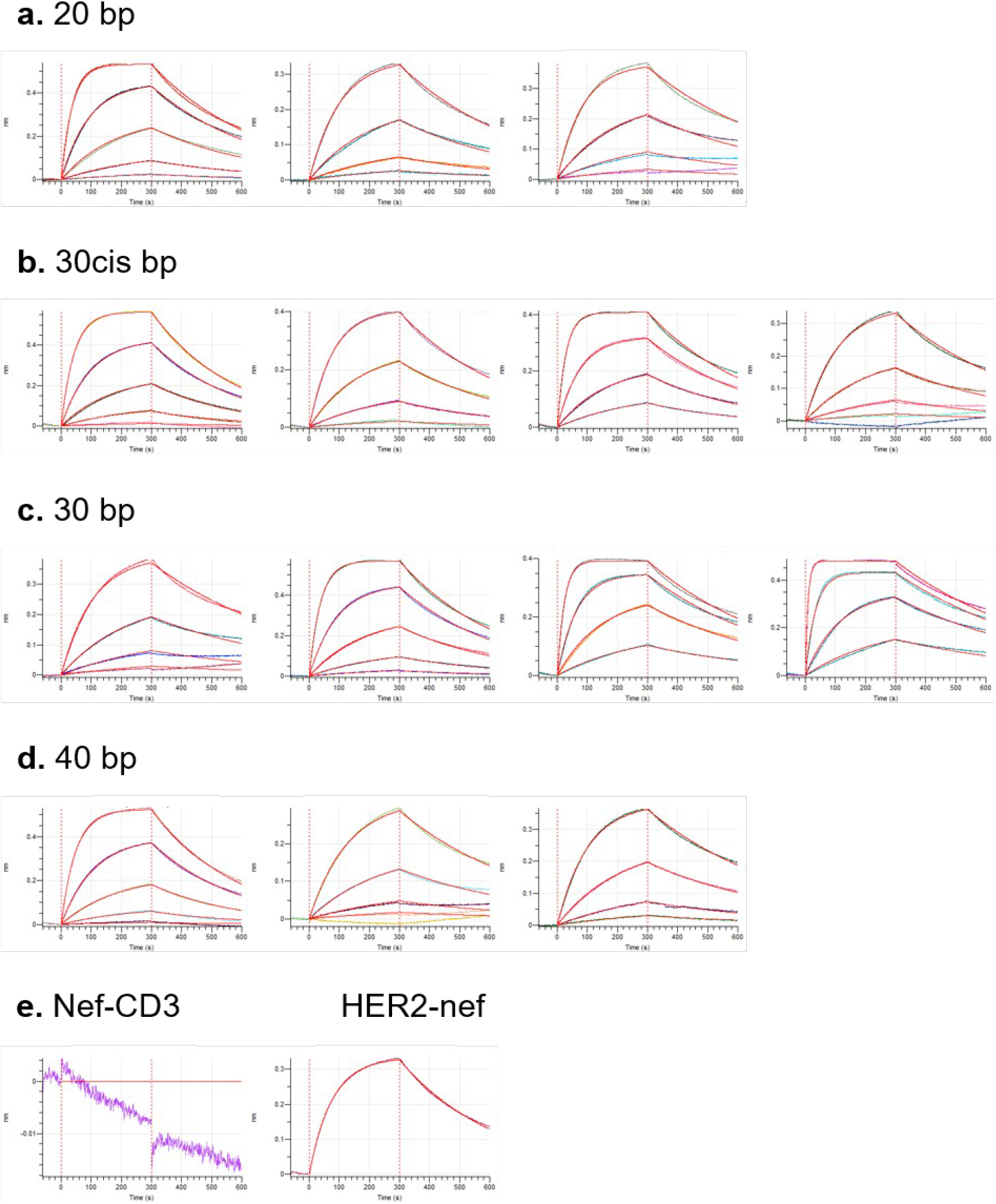
Kinetic analysis of DNab binding to HER2 by Biolayer interferometry (BLI). (a) DNab 20, (b) DNab 30cis, (c)DNab 30, (d) DNab 40, (e) Nef-CD3 and HER2-Nef. Binding kinetics were assessed by BLI using biotinylated recombinant HER2. Association and dissociation sensorgrams at four concentrations (1.8 to 150 nM). Experimental data (colored lines) are overlaid with the 1:1 global fit model (red lines). Data represent individual experimental replicates for each DNab.

**Extended Data Figure 5.**
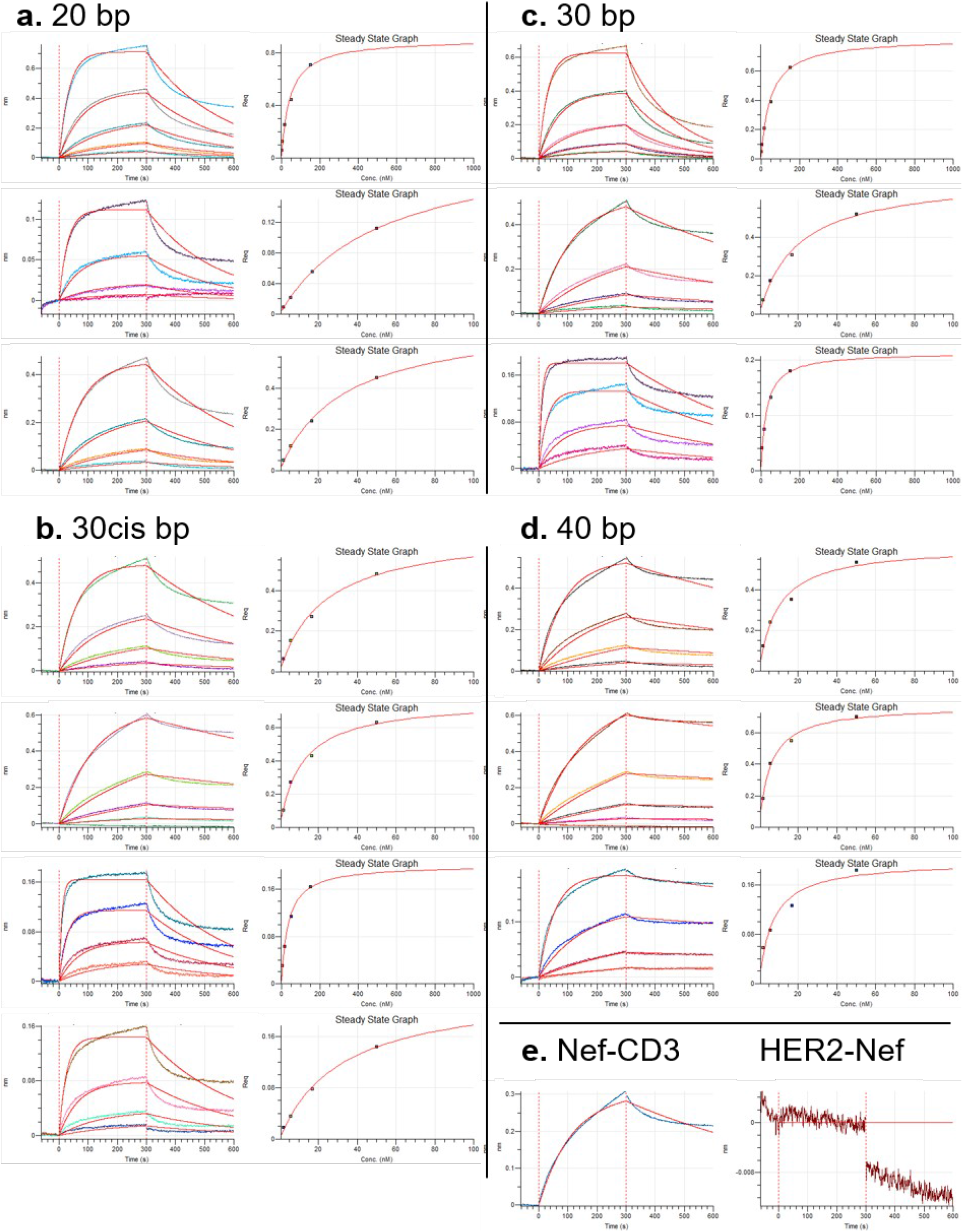
Kinetic analysis of DNab binding to CD3 measured by BLI. **(a)** DNab 20, **(b)** DNab 30cis, **(c)** DNa 30, **(d)** DNab 40, **(e)** Nef-CD3 and HER2-Nef. Binding kinetics were assessed by BLI using biotinylated recombinant CD3ε-δ heterodimer. **Left panels:** Association and dissociation curves (colored lines: experimental data at four concentrations ranging from 1.8 to 50 nM; red lines: 1:1 global fit model). **Right panels:** Steady-state affinity curves. n ≥ 3 independent experiments. Data represent individual experimental replicates for each DNab.

**Extended Data Figure 6.**
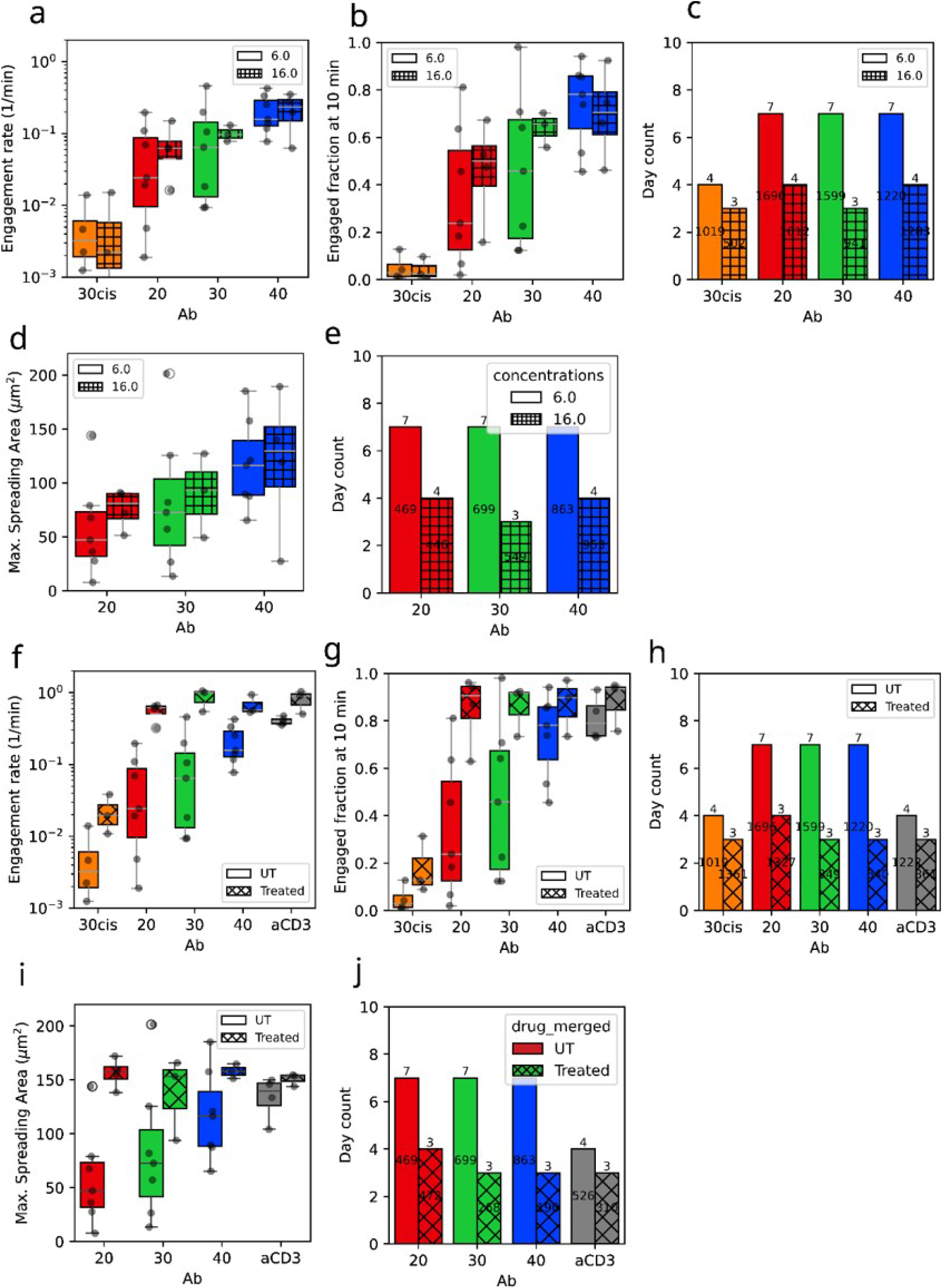
Effects of DNab concentration and neuraminidase treatment on T-cell spreading in the absence of ICAM-1. **(a,b)** Engagement rate and engaged fraction measured for all DNab formats at 6 nM and 16 nM. **(c)** Number of independent experiments and total number of analyzed cells for the datasets shown in (a,b). **(d)** Maximum spreading area of engaged cells for DNab at 6 nM and 16 nM. Analysis of 30cis was not shown because of the limited number of engagement events. **(e)** Number of independent experiments and total number of analyzed cells for the datasets shown in (d). **(f,g)** Engagement rate and engaged fraction measured for untreated (UT) and neuraminidase-treated cells at 6 nM DNab concentrations. **(h)** Number of independent experiments and total number of analysed cells for the datasets shown in (f,g). **(i)** Maximum spreading area of engaged cells untreated or treated with neuraminidase **(j)** Number of independent experiments and total number of analyzed cells for the dataset shown in (i).

**Extended Data Figure 7.**
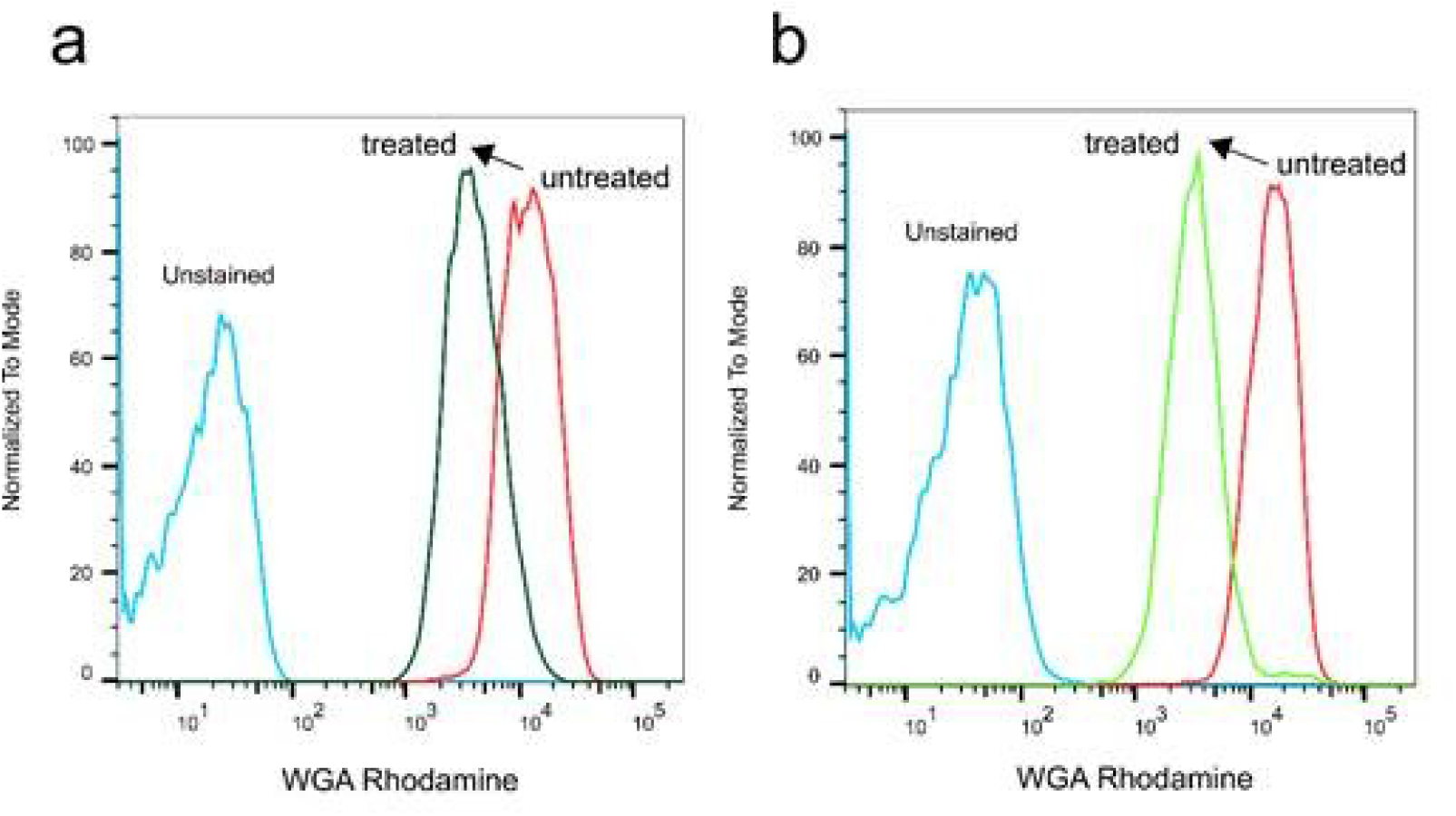
Effect of Neuraminidase treatment on T cell by flow cytometry. Representative flow cytometry histogram from 2 independent experiments **(a,b)** showed reduction in WGA-Rhodamine fluorescence after neuraminidase treatment compared with untreated cells.

**Extended Data Figure 8.**
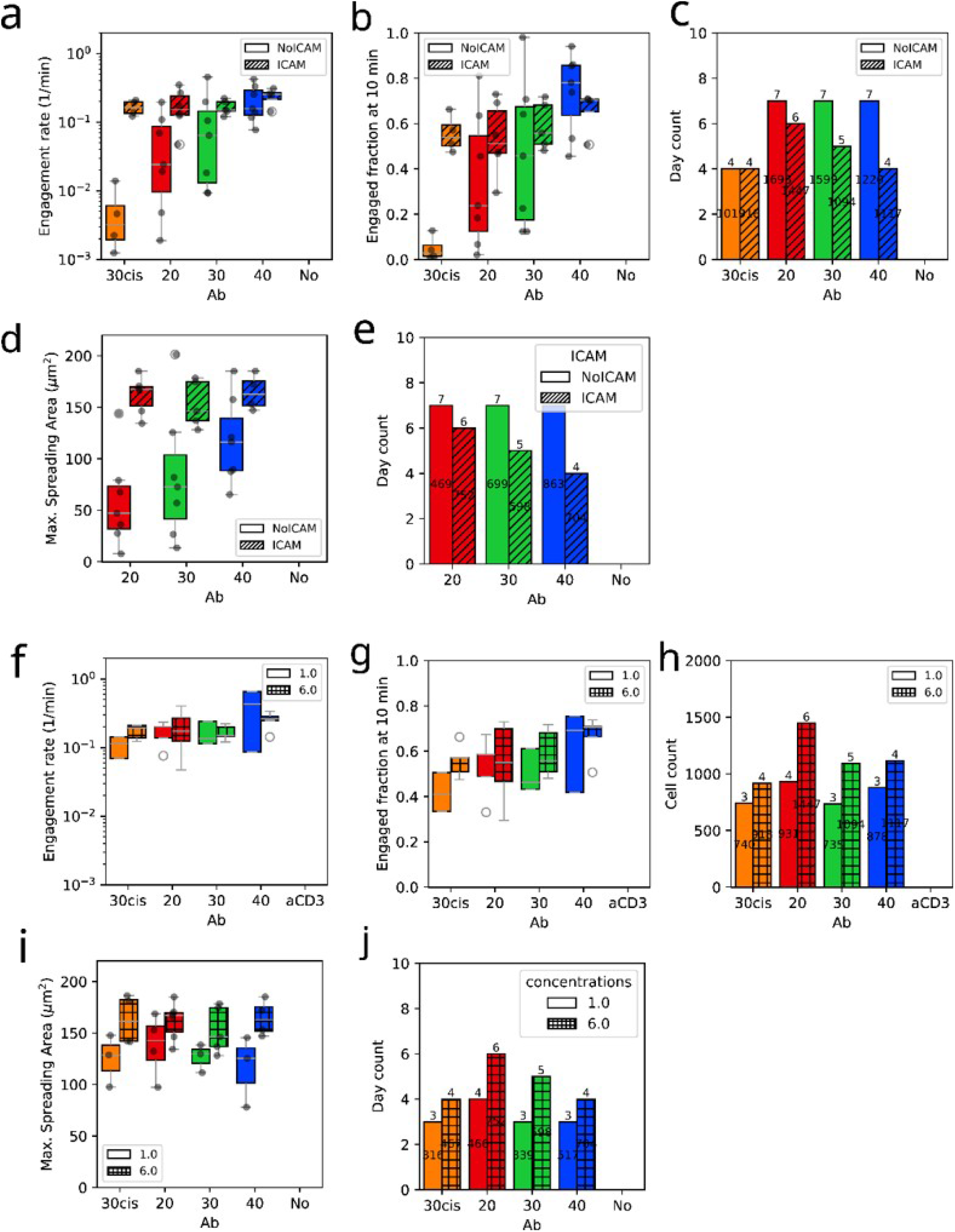
Effects of ICAM-1 and DNAb concentration on T-cell spreading. **(a,b)** Engagement rate and engaged fraction measured for the DNAb formats at 6 nM concentration in the absence or presence of ICAM-1. **(c)** Number of independent experiments and total number of analyzed cells for the datasets shown in (a,b). **(d)** Maximum spreading area of engaged cells for DNAb formats in the absence or presence of ICAM-1. **(e)** Number of independent experiments and total number of analyzed cells for the datasets shown in (d). **(f,g)** Engagement rate and engaged fraction measured for the indicated DNAb formats at 1 nM and 6 nM in the presence of ICAM-1. **(h)** Number of independent experiments and total number of analyzed cells for the datasets shown in (f,g). **(i)** Maximum spreading area of engaged cells for the indicated DNAb formats at 1 nM and 6 nM in the presence of ICAM-1. **(j)** Number of independent experiments and total number of analyzed cells for the datasets shown in (i).

**Extended Data Figure 9.**
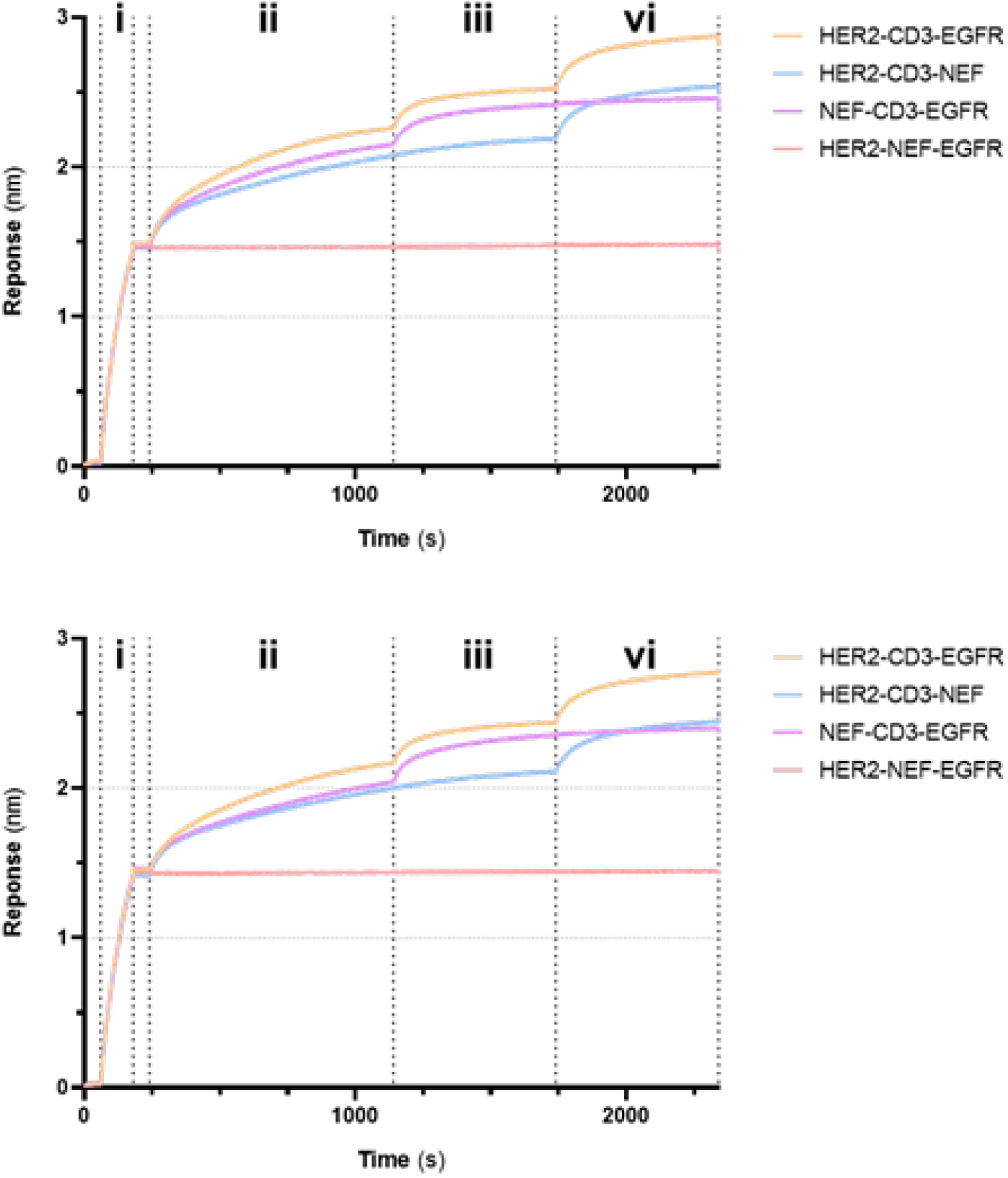
Trispecific TCE enables simultaneous engagement of three distinct target epitopes. BLI sensorgrams demonstrating the multispecificity of the DNabs (i) Biotinylated CD3ε-δ (1.25 µg/mL) was immobilized on streptavidin biosensors. (ii) Dipping of sensors into new wells containing EGFR-Fc (20 µg/mL) + DNabs, followed by (iv) transfer to wells containing a mixture of HER2-Fc (20 µg/mL), EGFR-Fc, and DNabs. Data show two additional independent experimental replicates for each construct to demonstrate the reproducibility of the multi-step binding profile.

## References

1. Fenis, A., Demaria, O., Gauthier, L., Vivier, E. & Narni-Mancinelli, E. New immune cell engagers for cancer immunotherapy. Nat Rev Immunol 24, 471–486 (2024).

2. Dustin, M. L. The immunological synapse. Cancer Immunol Res 2, 1023–1033 (2014).

3. Shimabukuro-Vornhagen, A. et al. Cytokine release syndrome. J Immunother Cancer 6, 56 (2018).

4. Dang, K. et al. Attenuating CD3 affinity in a PSMAxCD3 bispecific antibody enables killing of prostate tumor cells with reduced cytokine release. J Immunother Cancer 9, e002488 (2021).

5. Shah, D., Soper, B. & Shopland, L. Cytokine release syndrome and cancer immunotherapies – historical challenges and promising futures. Front Immunol 14, 1190379 (2023).

6. Belardi, B., Son, S., Felce, J. H., Dustin, M. L. & Fletcher, D. A. Cell-cell interfaces as specialized compartments directing cell function. Nat Rev Mol Cell Biol 21, 750–764 (2020).

7. Roda-Navarro, P. & AJ lvarez-Vallina, L. Understanding the Spatial Topology of Artificial Immunological Synapses Assembled in T Cell-Redirecting Strategies: A Major Issue in Cancer Immunotherapy. Frontiers in Cell and Developmental Biology 7, 370 (2020).

8. Zhao, L. et al. A novel CD19/CD22/CD3 trispecific antibody enhances therapeutic efficacy and overcomes immune escape against B-ALL. Blood 140, 1790–1802 (2022).

9. Tapia-Galisteo, A. et al. Trispecific T-cell engagers for dual tumor-targeting of colorectal cancer. Oncoimmunology 11, 2034355 (2022).

10. Zhou, X. et al. Using protein geometry to optimize cytotoxicity and the cytokine window of a ROR1 specific T cell engager. Frontiers in Immunology 15, 1323049 (2024).

11. Leithner, A. et al. Solution structure and synaptic analyses reveal determinants of bispecific T cell engager potency. Proc Natl Acad Sci U S A 122, e2425781122.

12. Chen, Y. et al. A Self-Assembled Protein Platform for Plug-and-Play Customization of Multivalent Artificial Antibodies and Antibody-Drug Conjugates. Angew Chem Int Ed 64, e202506426 (2025).

13. Lin, F. et al. Multimodal targeting chimeras enable integrated immunotherapy leveraging tumor-immune microenvironment. Cell 187, 7470–7491.e32 (2024).

14. Messaoudi, S. et al. Rapid Systematic Screening of Bispecific Antibody Surrogate Geometries for T-Cell Engagement Using DNA Nanotechnology. Journal of the American Chemical Society 146, 29824–29835 (2024).

15. Tang, R. et al. A Chimeric Conjugate of Antibody and Programmable DNA Nanoassembly Smartly Activates T Cells for Precise Cancer Cell Targeting. Angew Chem Int Ed 61, (2022).

16. Li, H.-D. et al. A Spatially Controllable DNA Origami-Based T-Cell Cluster Engager for Precise Tumor Elimination. J. Am. Chem. Soc. (2026).

17. Bannas, P., Hambach, J. & Koch-Nolte, F. Nanobodies and Nanobody-Based Human Heavy Chain Antibodies As Antitumor Therapeutics. Front. Immunol. 8, (2017).

18. Díaz-Bello, B. et al. Quantitative Microscopy for Cell–Surface and Cell–Cell Interactions in Immunology. Bio-Protocol 15, (2025).

19. Torro, R. et al. Celldetective: an AI-enhanced image analysis tool for unraveling dynamic cell interactions. eLife https://doi.org/10.7554/eLife.105302.1 (2025) doi:10.7554/eLife.105302.1.

20. Valignat, M.-P., Theodoly, O., Gucciardi, A., Hogg, N. & Lellouch, A. C. T Lymphocytes Orient against the Direction of Fluid Flow during LFA-1-Mediated Migration. Biophysical Journal 104, 322–331 (2013).

21. Davis, S. J. & van der Merwe, P. A. The kinetic-segregation model: TCR triggering and beyond. Nat Immunol 7, 803–809 (2006).

22. Dillard, P., Varma, R., Sengupta, K. & Limozin, L. Ligand-Mediated Friction Determines Morphodynamics of Spreading T Cells. Biophysical Journal 107, 2629–2638 (2014).

23. Sengupta, K., Dillard, P. & Limozin, L. Morphodynamics of T-lymphocytes: scanning to spreading. Biophysical Journal 123, 2224–2233 (2024).

24. Brodovitch, A., Bongrand, P. & Pierres, A. T Lymphocytes Sense Antigens within Seconds and Make a Decision within One Minute. The Journal of Immunology 191, 2064–2071 (2013).

25. Moreau, H. D. et al. Signal strength regulates antigen-mediated T-cell deceleration by distinct mechanisms to promote local exploration or arrest. Proceedings of the National Academy of Sciences 112, 12151–12156 (2015).

26. Mayya, V. et al. Durable Interactions of T Cells with T Cell Receptor Stimuli in the Absence of a Stable Immunological Synapse. Cell Rep 22, 340–349 (2018).

27. Wilhelm, K. B. et al. Height, but not binding epitope, affects the potency of synthetic TCR agonists. Biophysical Journal 120, 3869–3880 (2021).

28. Kamal, M. A. et al. Physics of Organelle Membrane Bridging via Cytosolic Tethers is Distinct From Cell Adhesion. Front. Phys. 9, 750539 (2022).

29. Burton, J. et al. Optimising CAR-T cell sensitivity by engineering matched extracellular sizes between CAR/antigen and CD2/CD58 adhesion complexes. 2025.01.06.631424 Preprint at 10.1101/2025.01.06.631424 (2025).

30. Fenz, S. F. et al. Membrane fluctuations mediate lateral interaction between cadherin bonds. Nature Physics 13, 906–913 (2017).

31. Pierres, A., Benoliel, A.-M., Touchard, D. & Bongrand, P. How Cells Tiptoe on Adhesive Surfaces before Sticking. Biophysical Journal 94, 4114–4122 (2008).

32. GonzaS lez Gutierrez, C., et al. Decoupling Individual Host Response and Immune Cell Engager Cytotoxic Potency. ACS Nano 19, 2089–2098 (2025).

33. Figenschau, S. L. et al. ICAM-11 expression is induced by proinflammatory cytokines and associated with TLS formation in aggressive breast cancer subtypes. Sci Rep 8, 11720 (2018).

34. Park, S. et al. Immunoengineering can overcome the glycocalyx armour of cancer cells. Nat. Mater. 23, 429–438 (2024).

35. Xiao, H., Woods, E. C., Vukojicic, P. & Bertozzi, C. R. Precision glycocalyx editing as a strategy for cancer immunotherapy. Proceedings of the National Academy of Sciences 113, 10304–10309 (2016).

36. Yang, Z. et al. Targeted desialylation and cytolysis of tumour cells by fusing a sialidase to a bispecific T-cell engager. *Nat*. Biomed. Eng 8, 499–512 (2024).

37. Choudhuri, K., Wiseman, D., Brown, M. H., Gould, K. & van der Merwe, P. A. T-cell receptor triggering is critically dependent on the dimensions of its peptide-MHC ligand. Nature 436, 578–582 (2005).

38. Xiao, Q., et al. Size-dependent activation of CAR-T cells. Sci Immunol 7, eabl3995 (2022).

39. Faroudi, M. et al. Lytic versus stimulatory synapse in cytotoxic T lymphocyte/target cell interaction: Manifestation of a dual activation threshold. Proceedings of the National Academy of Sciences 100, 14145–14150 (2003).

40. Nolan-Stevaux, O. & Smith, R. Logic-gated and contextual control of immunotherapy for solid tumors: contrasting multi-specific T cell engagers and CAR-T cell therapies. Frontiers in Immunology 15, 1490911 (2024).

41. Bergamaschi, C. et al. Innovative strategies for T cell engagers for cancer immunotherapy. MAbs 17, 2531223.

42. Bouchet, J. et al. Inhibition of the Nef regulatory protein of HIV-1 by a single-domain antibody. Blood 117, 3559–3568 (2011).

43. Nevoltris, D. et al. Conformational Nanobodies Reveal Tethered Epidermal Growth Factor Receptor Involved in EGFR/ErbB2 Predimers. ACS Nano 9, 1388–1399 (2015).

44. Behar, G. et al. Isolation and characterization of anti-FcgammaRIII (CD16) llama single-domain antibodies that activate natural killer cells. Protein engineering, design & selection : PEDS 21, 1–10 (2008).

45. Even-Desrumeaux, K. et al. Masked Selection: A Straightforward and Flexible Approach for the Selection of Binders Against Specific Epitopes and Differentially Expressed Proteins by Phage Display. Molecular & Cellular Proteomics : MCP 13, 653 (2013).

