## Supplementary information for "Tuning T-cell immunological synapse by modular DNA-Nanobody engagers for precision immunotherapy"

+ Equal contribution. * Equal contribution

### TABLE OF CONTENTS

**Supplementary Figures 1-20**

**Supplementary Tables 1-3**

**Supplementary Videos 1-2**

**Supplementary Figure 1.** Schematic representation and size estimation of DNA–Nanobody (DNAb) constructs.

**Supplementary Figure 2.** Pairwise estimation analysis of engagement rate (6 nM DNAbs, without ICAM).

**Supplementary Figure 3.** Pairwise estimation analysis of engagement rate (16 nM DNAbs, without ICAM).

**Supplementary Figure 4.** Pairwise estimation analysis of maximum spreading area (6 nM DNAbs, without ICAM).

**Supplementary Figure 5.** Pairwise estimation analysis of maximum spreading area (16 nM DNAbs, without ICAM).

**Supplementary Figure 6.** Pairwise estimation analysis of neuraminidase-mediated changes in engagement rate (6 nM DNAbs, without ICAM).

**Supplementary Figure 7.** Pairwise estimation analysis of engagement rate following neuraminidase treatment (16 nM DNAbs, without ICAM).

**Supplementary Figure 8.** Pairwise estimation analysis of engagement rate (6 nM DNAbs, with ICAM-1).

**Supplementary Figure 9.** Pairwise estimation analysis of spreading area (6 nM DNAbs, with ICAM-1).

**Supplementary Figure 10.** Pairwise estimation analysis of stop rate (6 nM DNAbs, with ICAM-1).

**Supplementary Figure 11.** Pairwise estimation analysis of stop rate (1 nM DNAbs, with ICAM-1).

**Supplementary Figure 12.** Pairwise estimation analysis of stop fraction (1 nM DNAbs, with ICAM-1).

**Supplementary Figure 13.** Pairwise estimation analysis of CD45 exclusion.

**Supplementary Figure 14.** Pairwise estimation analysis of cytotoxicity (lysis EC50 and maximal lysis).

**Supplementary Figure 15.** Pairwise estimation analysis of IFN-γ secretion (EC50 and maximal response).

**Supplementary Figure 16.** RICM image normalization and background validation.

**Supplementary Figure 17.** Cell segmentation, tracking and trajectory reconstruction workflow.

**Supplementary Figure 18.** Determination of spreading onset from RICM contact area.

**Supplementary Figure 19.** Determination of arrest onset from cell velocity analysis.

**Supplementary Figure 20.** Quantification of CD45 exclusion by radial intensity analysis.

**Supplementary Table 1.** Binding parameters of DNabs determined by flow cytometry.

**Supplementary Table 2.** Survival of MCF-7-HER2$^{+}$ target cells after 4 h of DNab-mediated cytotoxicity.

**Supplementary Table 3.** Quantification of HER2 and EGFR surface expression on MCF-7, MCF-7-HER2$^{+}$, and A431 cell lines.

**Supplementary Video 1.** T cell engagement on HER2-coated surfaces in the presence of different length of DNAb at 6nM.

**Supplementary Video 2.** T cell arrest on HER2- and ICAM-coated surfaces in presence of DNAb at different concentration.

**Supplementary Video 3.** T cell-mediated killing of MCF-7-HER2$^{+}$ target cells in the presence of different length of DNAb

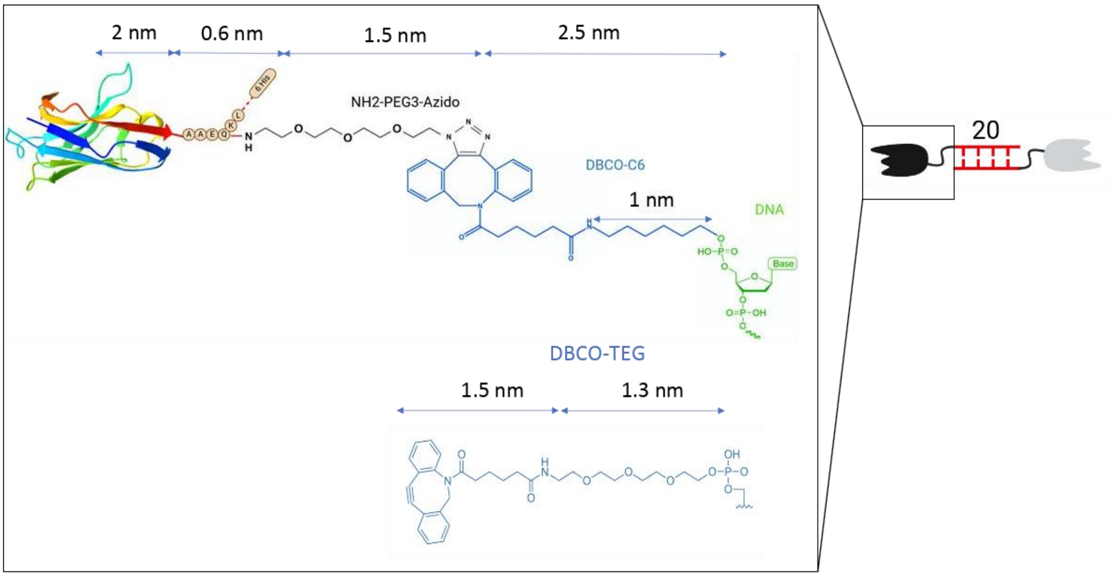


1 pb = 0.34 nm

30cis: 2+4.6+2+4.9+2 = 15.5 -> **16 nm**

20: 2+4.6+0.34x20+4.6+2 = 20.0 -> **20 nm**

30 : 2+4.9+0.34x30+4.9+2 = 24.0 -> **24 nm**

40 : 2+4.6+0.34x40+4.6+2 = 26.8 -> **27 nm**

**Supplementary Figure 1 | Schematic representation of DNAb size estimation. (a)** Each DNAb construct is composed of two nanobodies positioned at opposite extremities, with the antigen-binding epitopes located approximately 2 nm from the C-terminus of each nanobody. The nanobodies carry a C-terminal c-Myc tag, which enables site-specific conjugation to the linker (NH2 -PEG3 -azide) via transglutaminase-mediated coupling between the glutamine residue within the c-Myc tag and the primary amine of the linker, resulting in a stable covalent amide bond. The azide-functionalized linker subsequently reacts with a DBCO-modified oligonucleotide through strain-promoted azide–alkyne cycloaddition (SPAAC). The dimensions of each molecular component are indicated in the schematic.Two types of DBCO-functionalized oligonucleotides were used: one comprising a DBCO moiety followed by a six-carbon spacer (C6), and another incorporating a DBCO–TEG linker. **(b)** The overall size of the DNAb constructs was calculated based on the dimensions of these individual components.

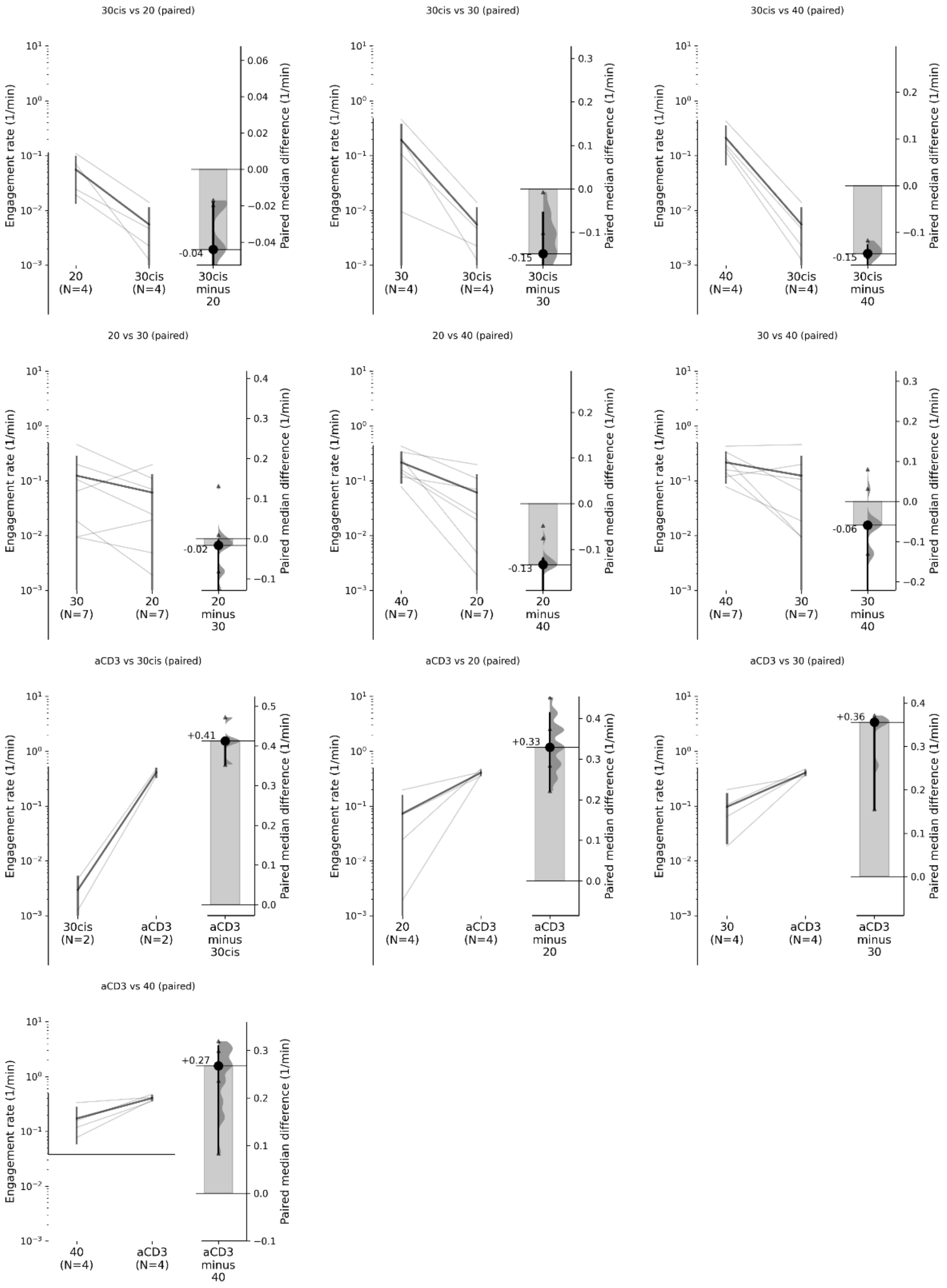


**Supplementary Figure 2.** Pairwise estimation analysis of Engagement rate for DNAbs at a concentration of 6 nM in the absence of ICAM, corresponding to the plot shown in Fig. 2d. The significance is indicated by # in Fig. 2d when the average difference and 95% confidence interval, shown as a black circle with error bar, do not cross the horizontal zero line, indicating reliable measured differences.

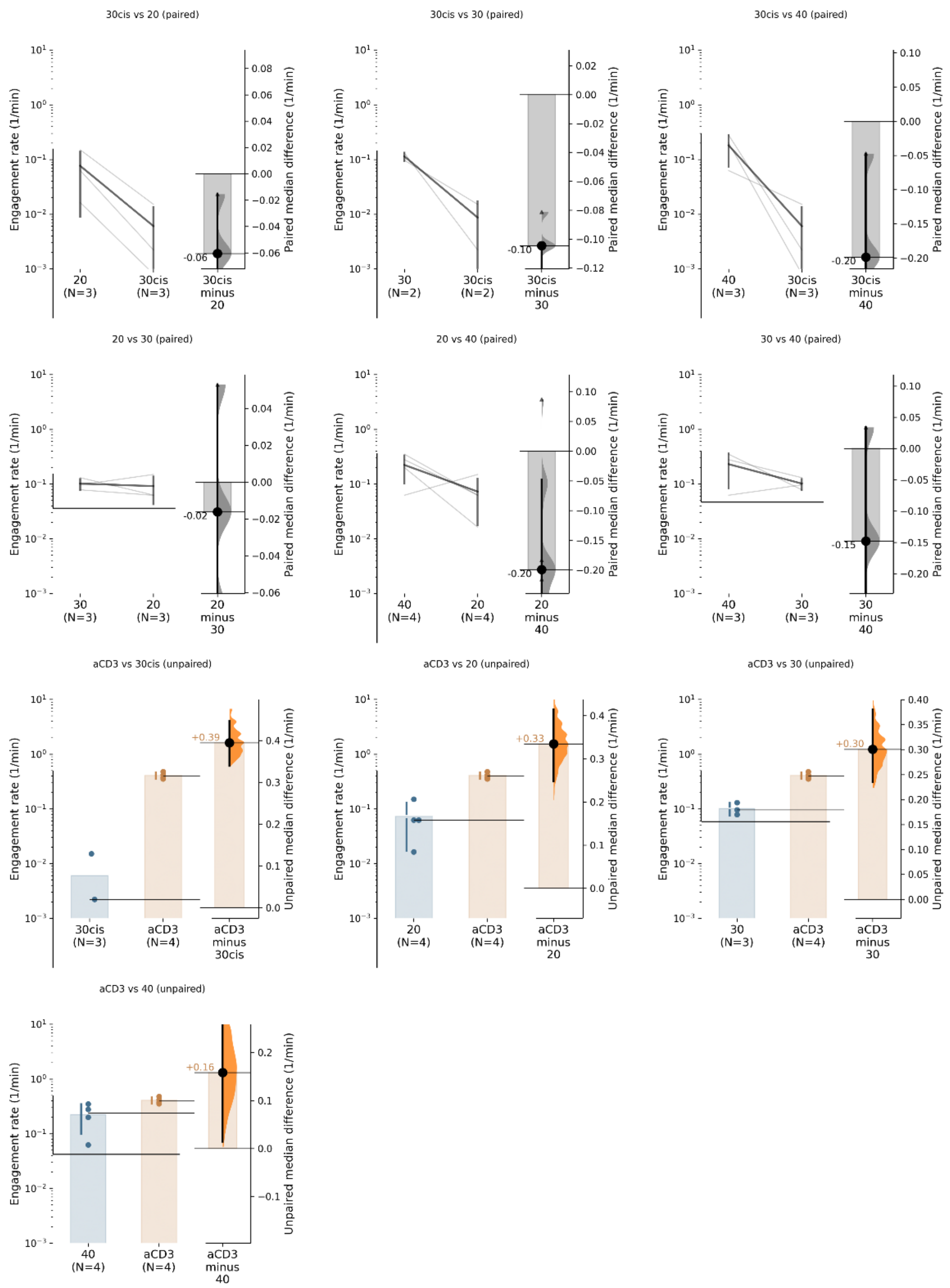


**Supplementary Figure 3.** Pairwise estimation analysis of engagement rate for DNAbs at a concentration of 16 nM in the absence of ICAM, corresponding to the plot shown in Fig. 2e. Comparisons were analyzed as paired when measurements were available on the same experimental dates, and as unpaired for comparisons with aCD3 when matched dates were not available. Significance is indicated by # in Fig. 2e when the average difference and 95% confidence interval, shown as a black circle with an error bar, do not cross the horizontal zero line, indicating reliable measured differences.

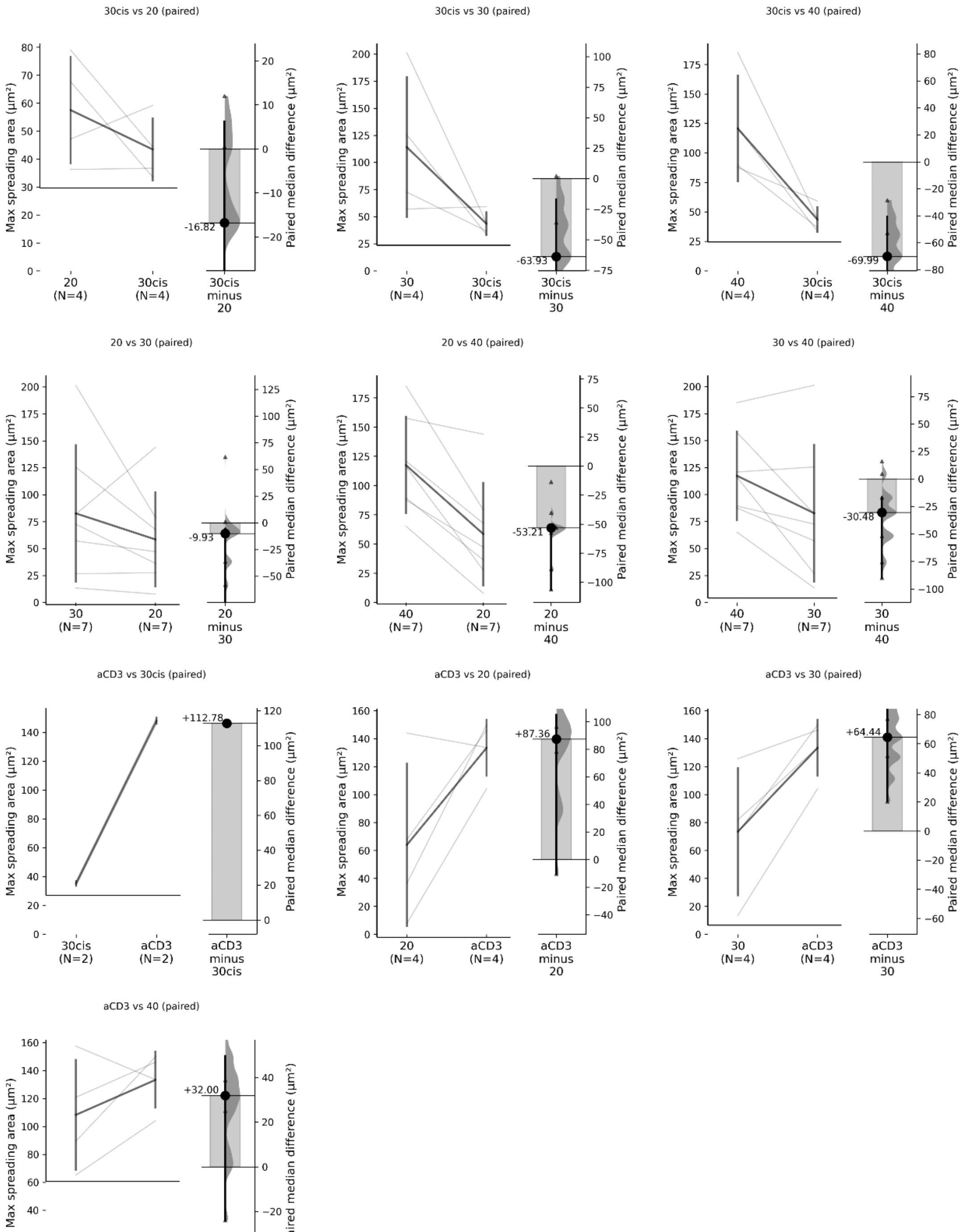


**Supplementary Figure 4.** Pairwise estimation analysis of Max spreading area for DNAbs at a concentration of 6 nM in the absence of ICAM, corresponding to the plot shown in Fig. 2g. Significance is indicated by # in Fig. 2g when the average difference and 95% confidence interval, shown as a black circle with an error bar, do not cross the horizontal zero line, indicating reliable measured differences.

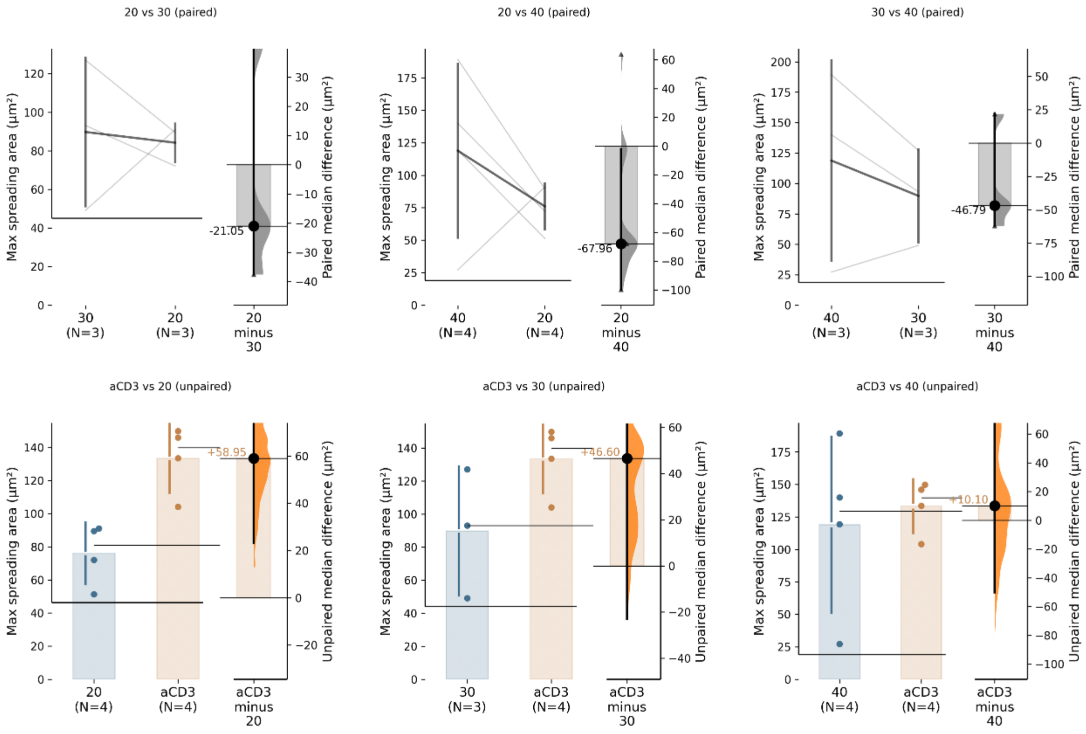


**Supplementary Figure 5.** Pairwise estimation analysis of Max spreading area for DNAbs at a concentration of 16 nM in the absence of ICAM, corresponding to the plot shown in Fig. 2h. Comparisons were analyzed as paired when measurements were available on the same experimental dates, and as unpaired for comparisons when matched dates were not available. Significance is indicated by # in Fig. 2h when the average difference and 95% confidence interval, shown as a black circle with an error bar, do not cross the horizontal zero line, indicating reliable measured differences.

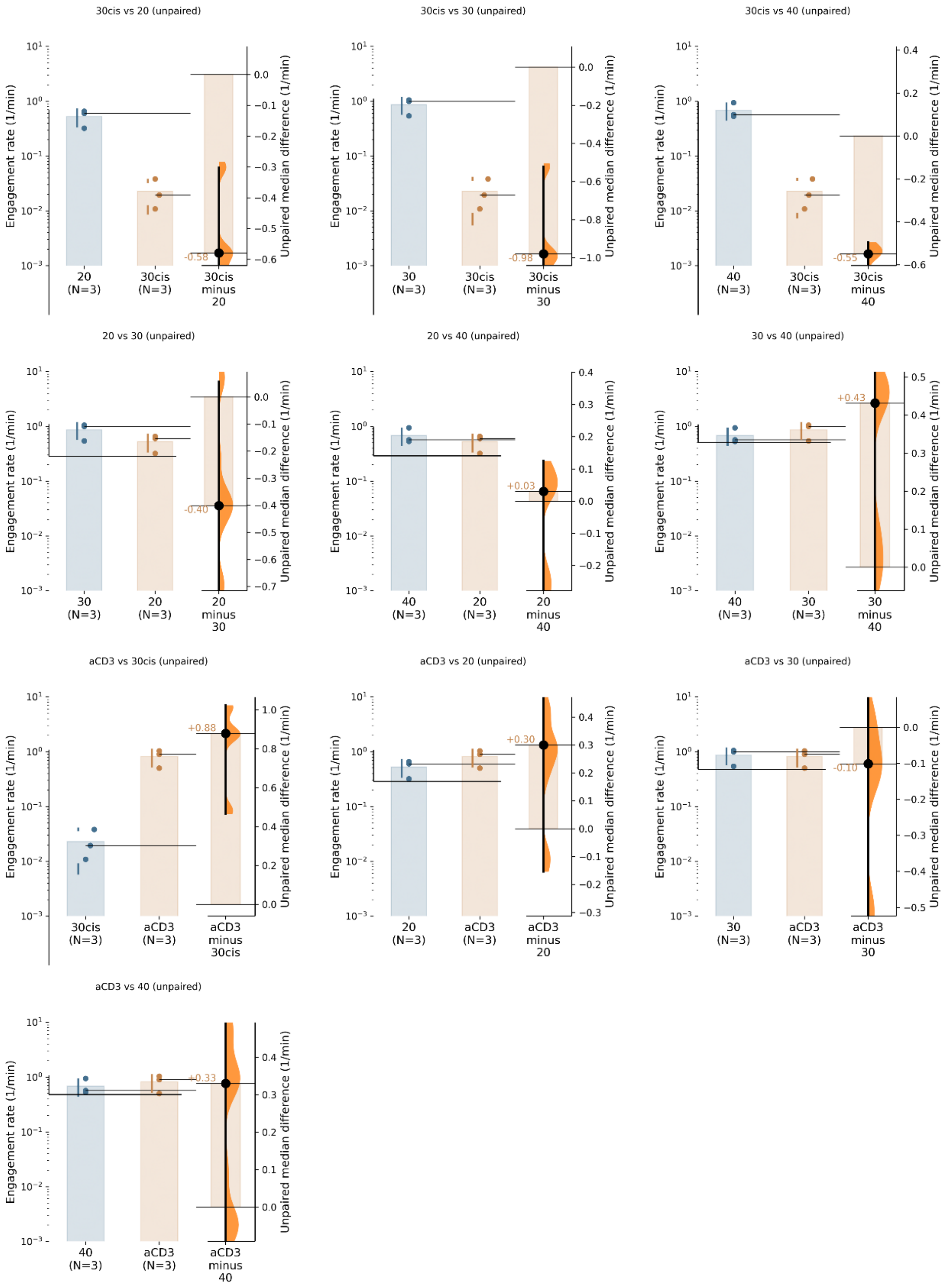


**Supplementary Figure 6.** Pairwise estimation analysis of effect of Neuraminidase on engagement rate for DNAbs at a concentration of 6 nM in the absence of ICAM, corresponding to the plot shown in Fig. 2j. Significance is indicated by # in Fig. 2j when the average difference and 95% confidence interval, shown as a black circle with an error bar, do not cross the horizontal zero line, indicating reliable measured differences.

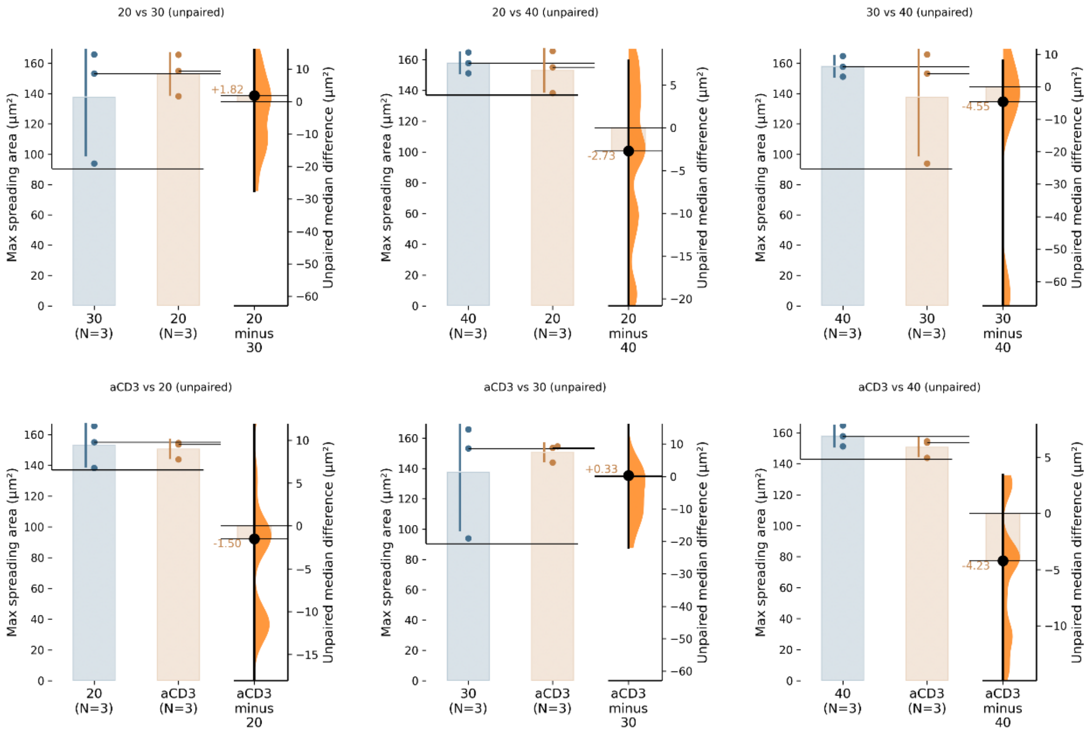


**Supplementary Figure 7.** Pairwise estimation analysis of engagement rate for DNAbs at a concentration of 16 nM in the absence of ICAM, corresponding to the plot shown in Fig. 2k. Significance is indicated by # in Fig. 2k when the average difference and 95% confidence interval, shown as a black circle with an error bar, do not cross the horizontal zero line, indicating reliable measured differences.

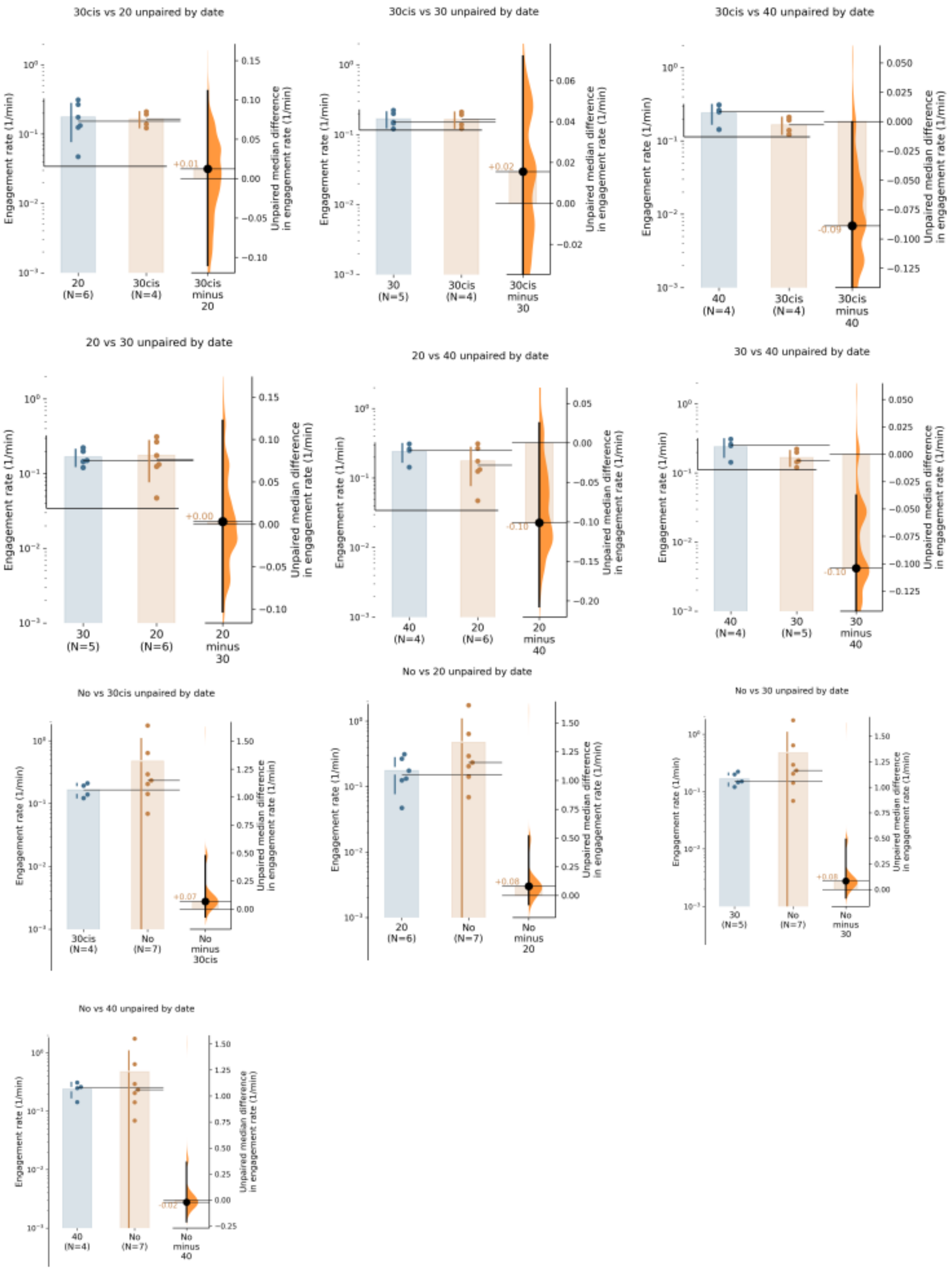


**Supplementary Figure 8.** Pairwise estimation analysis of Engagement rate for DNAbs at a concentration of 6 nM in the presence of ICAM, corresponding to the plot shown in Fig. 3c. The significance is indicated by # in Fig. 2c when the average difference and 95% confidence interval, shown as a black circle with error bar, do not cross the horizontal zero line, indicating reliable measured differences.

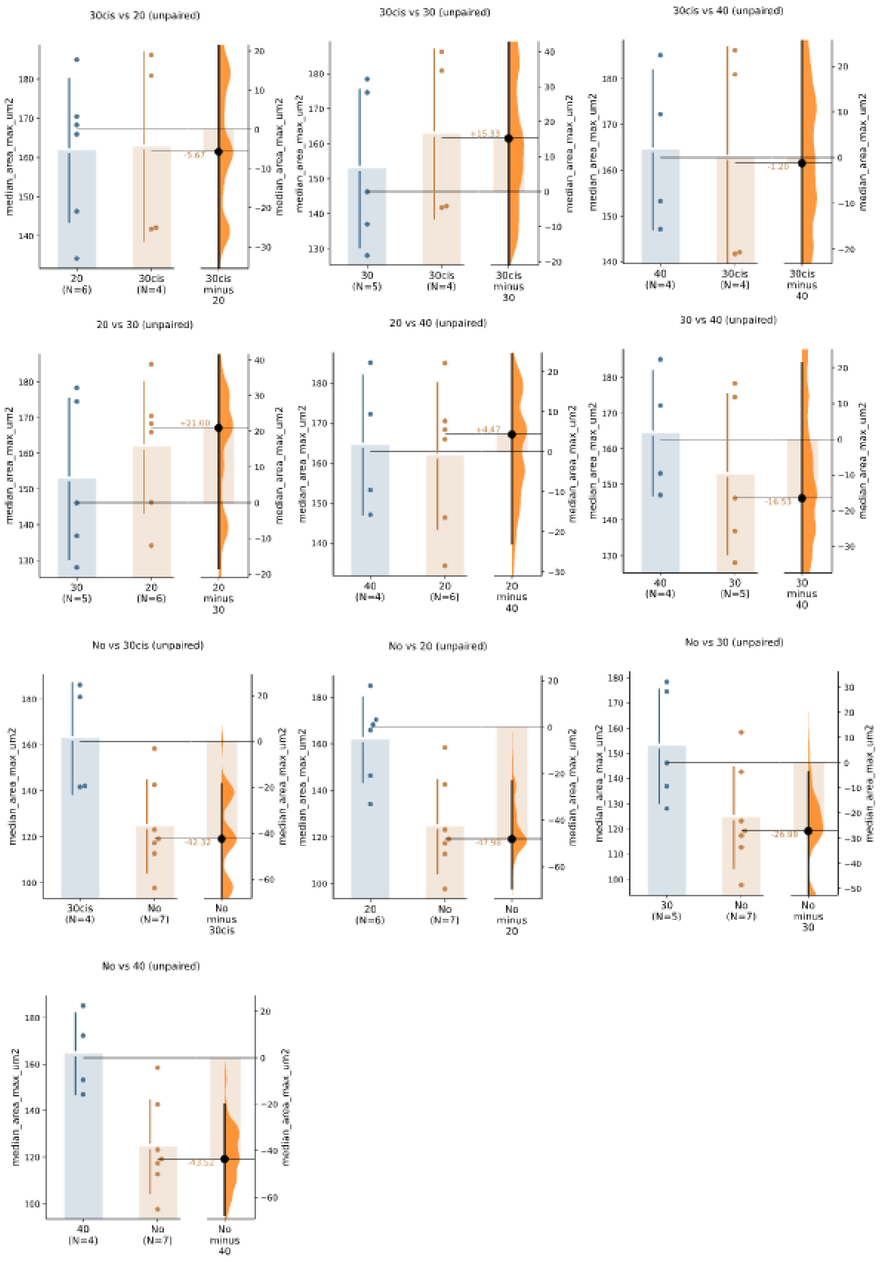


**Supplementary Figure 9.** Pairwise estimation analysis of Spreading area of DNAbs at a concentration of 6 nM in the presence of ICAM, corresponding to the plot shown in Fig. 3d. Significance is indicated by # in Fig. 3d when the average difference and 95% confidence interval, shown as a black circle with an error bar, do not cross the horizontal zero line, indicating reliable measured differences.

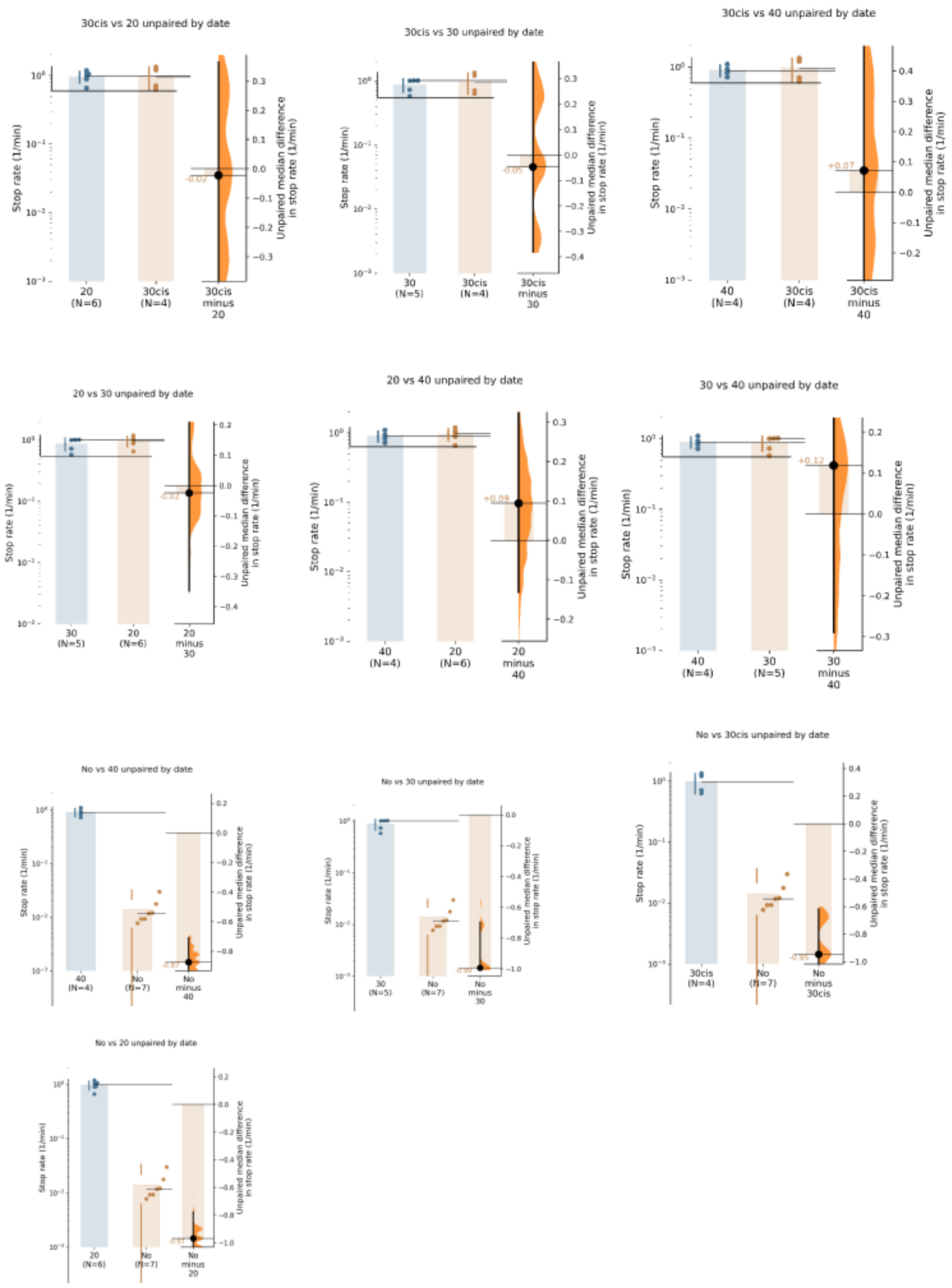


**Supplementary Figure 10.** Pairwise estimation analysis of Stop rate for DNAbs at a concentration of 6 nM in the presence of ICAM, corresponding to the plot shown in Fig. 3f. Significance is indicated by # in Fig. 3f when the average difference and 95% confidence interval, shown as a black circle with an error bar, do not cross the horizontal zero line, indicating reliable measured differences.

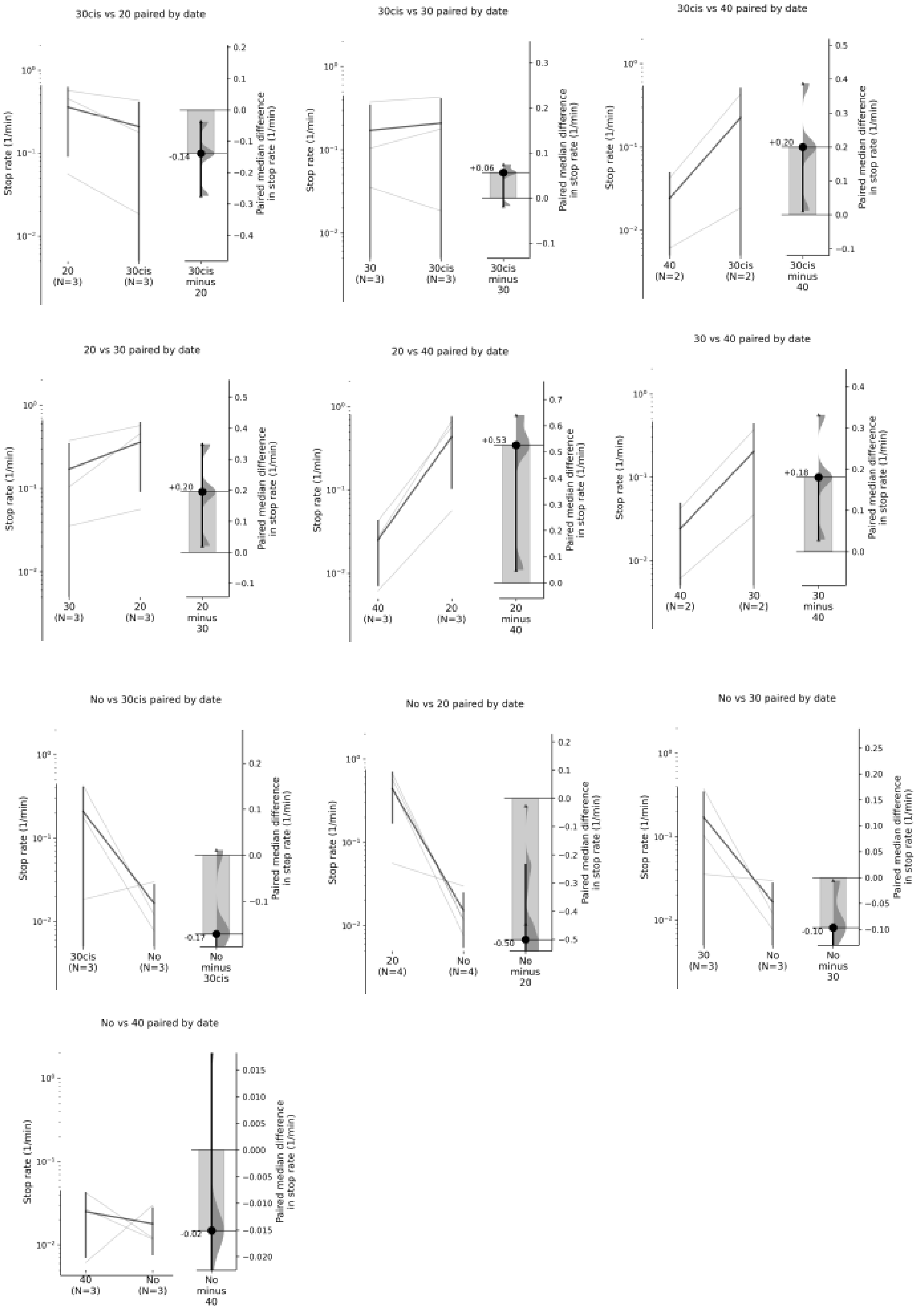


**Supplementary Figure 11.** Pairwise estimation analysis of Stop rate for DNAbs at a concentration of 1 nM in the presence of ICAM, corresponding to the plot shown in Fig. 3g. Significance is indicated by # in Fig. 3g when the average difference and 95% confidence interval, shown as a black circle with an error bar, do not cross the horizontal zero line, indicating reliable measured differences.

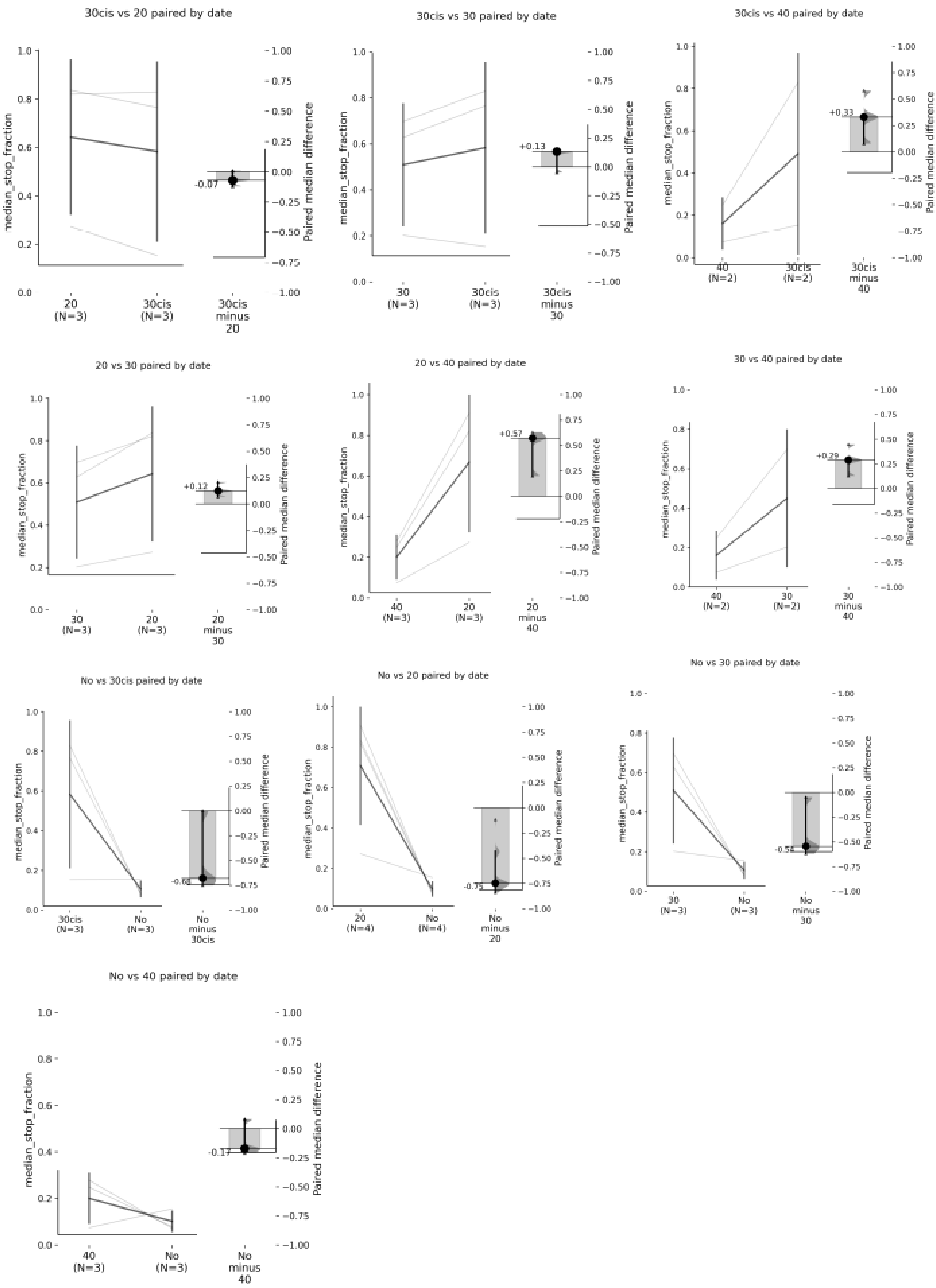


**Supplementary Figure 12.** Pairwise estimation analysis of Stop fraction for DNAbs at a concentration of 1 nM in the presence of ICAM, corresponding to the plot shown in Fig. 3h. Significance is indicated by # in Fig. 3h when the average difference and 95% confidence interval, shown as a black circle with an error bar, do not cross the horizontal zero line, indicating reliable measured differences.

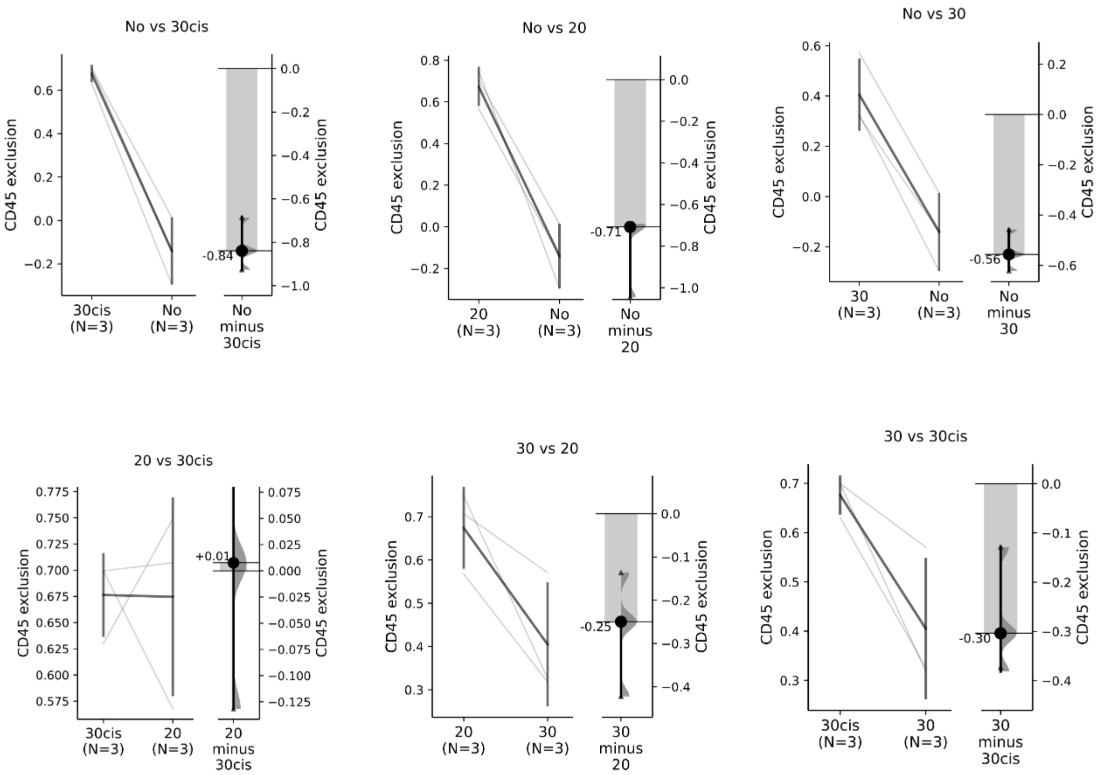


**Supplementary Figure 13.** Pairwise estimation analysis of Cd45 exclusion for DNAbs at a concentration of 6 nM in the presence of ICAM, corresponding to the plot shown in Fig. 3j. Significance is indicated by # in Fig. 3j when the average difference and 95% confidence interval, shown as a black circle with an error bar, do not cross the horizontal zero line, indicating reliable measured differences.

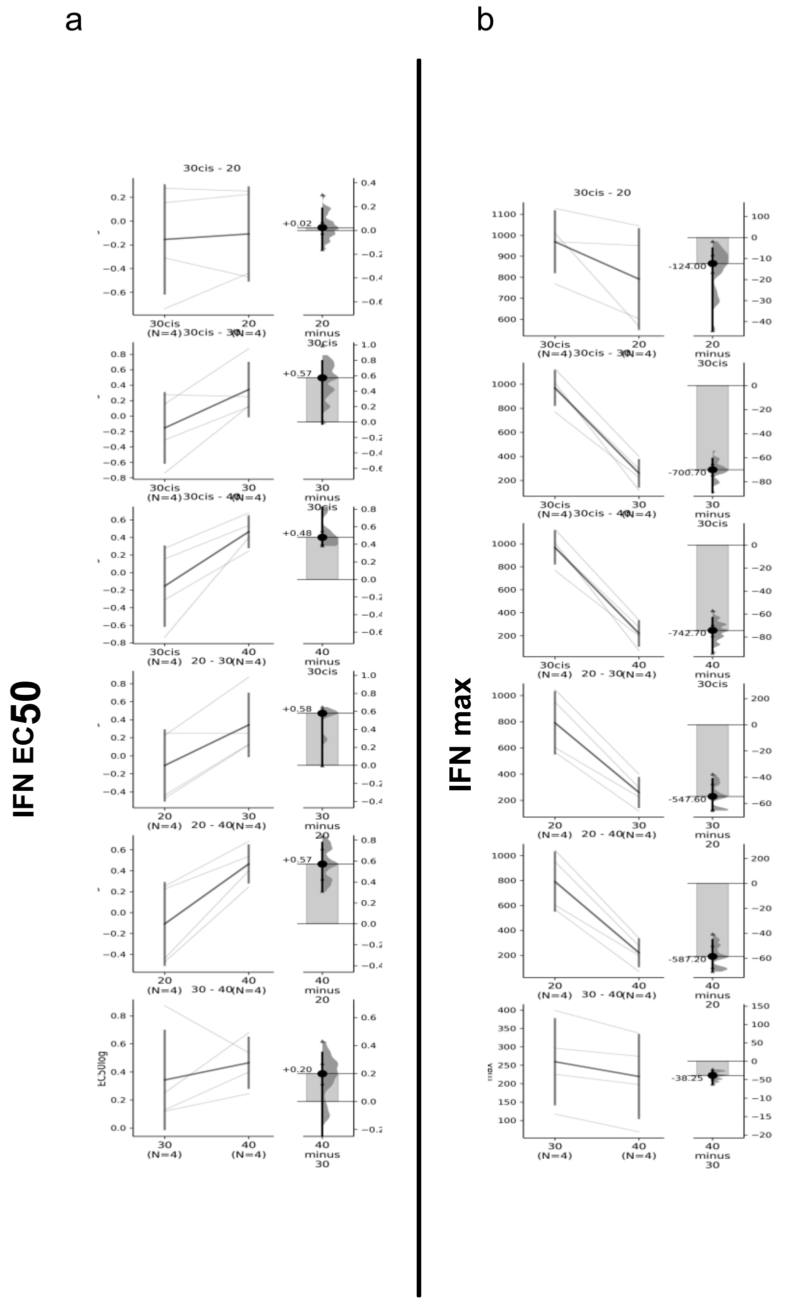


**Supplementary Figure 14.** Pairwise estimation analysis of lysis EC50 (**a**) and lysis maximum (**b**), corresponding to the plots shown in Fig. 4c and Fig. 4e, respectively.

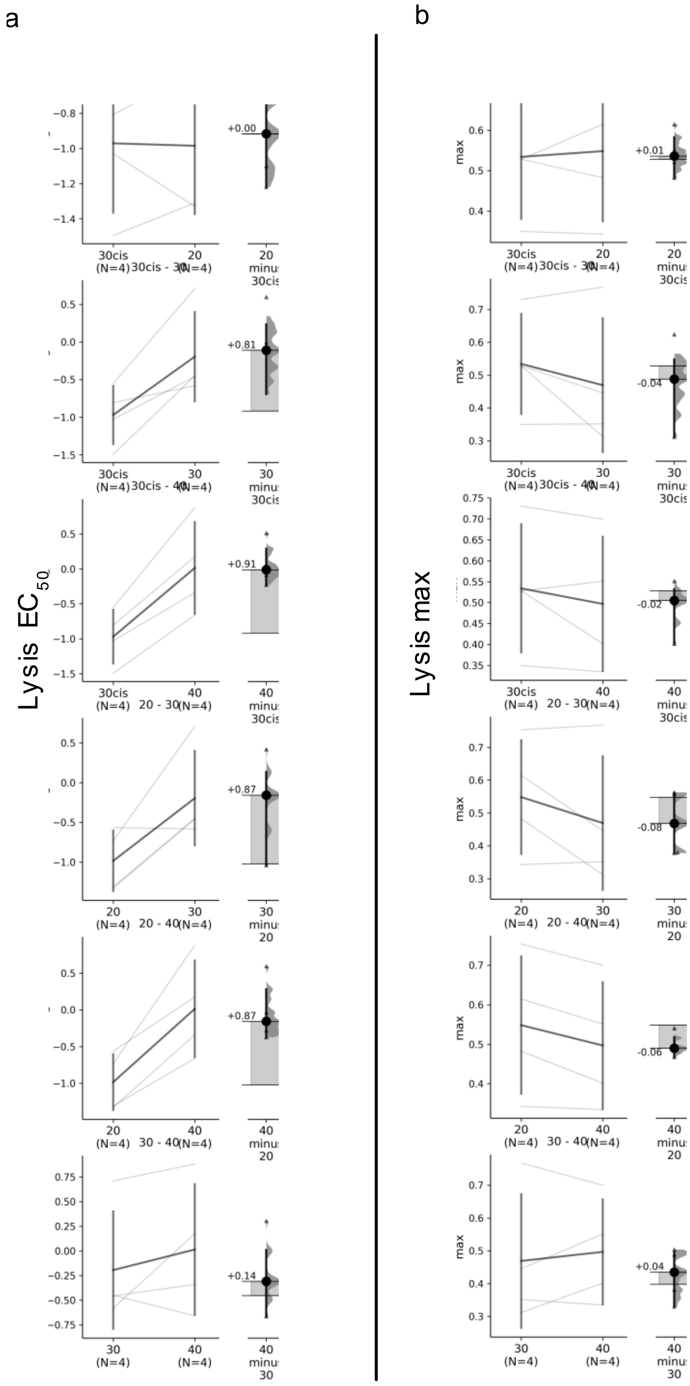


**Supplementary Figure 15.** Pairwise estimation analysis of IFN EC50 (**a**) and IFNmaximum (**b**), corresponding to the plots shown in Fig. 4d and Fig. 4f, respectively.

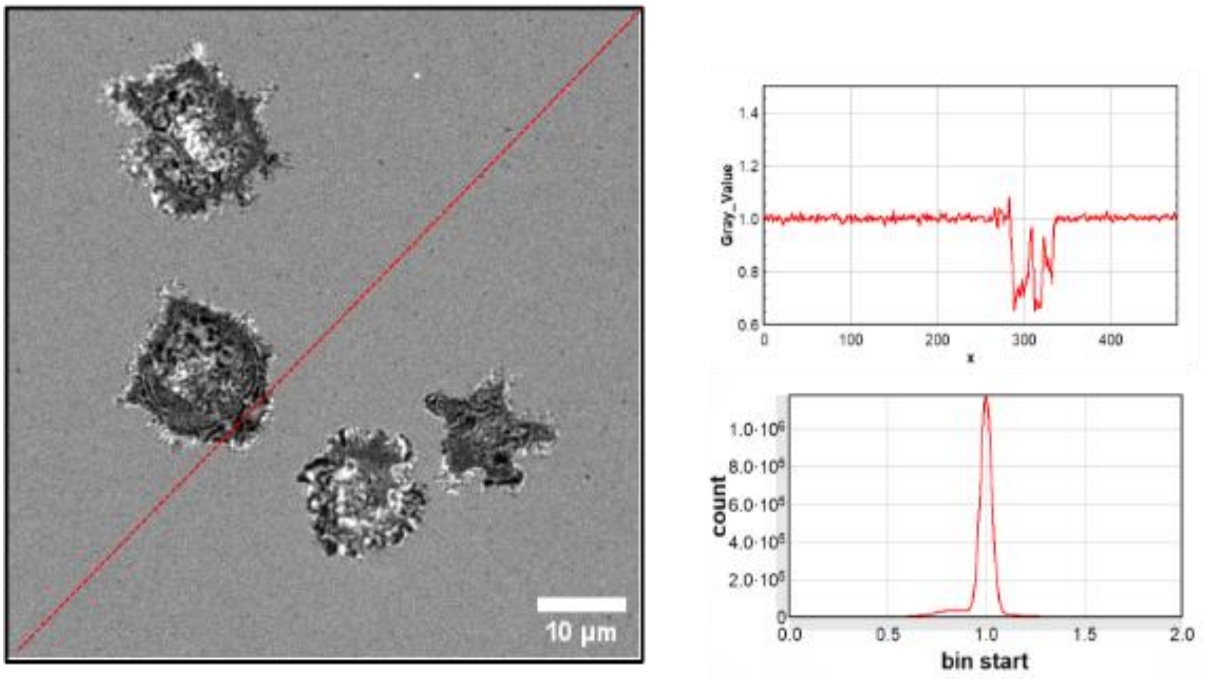


**Supplementary Figure 16. RICM normalization and background validation**. RICM images were normalized such that the background intensity was centered around 1. A representative RICM image is shown (left), with the red dashed line indicating the region used for intensity profiling. The corresponding line profile (top right) shows background values close to 1.0, while lower intensity values correspond to cell-adhered regions. To assess the quality of normalization, pixel intensities from all images across 60 frames were pooled into a single histogram (bottom right). This distribution exhibits a narrow peak centered at ~1.0, confirming consistent background normalization across the dataset.

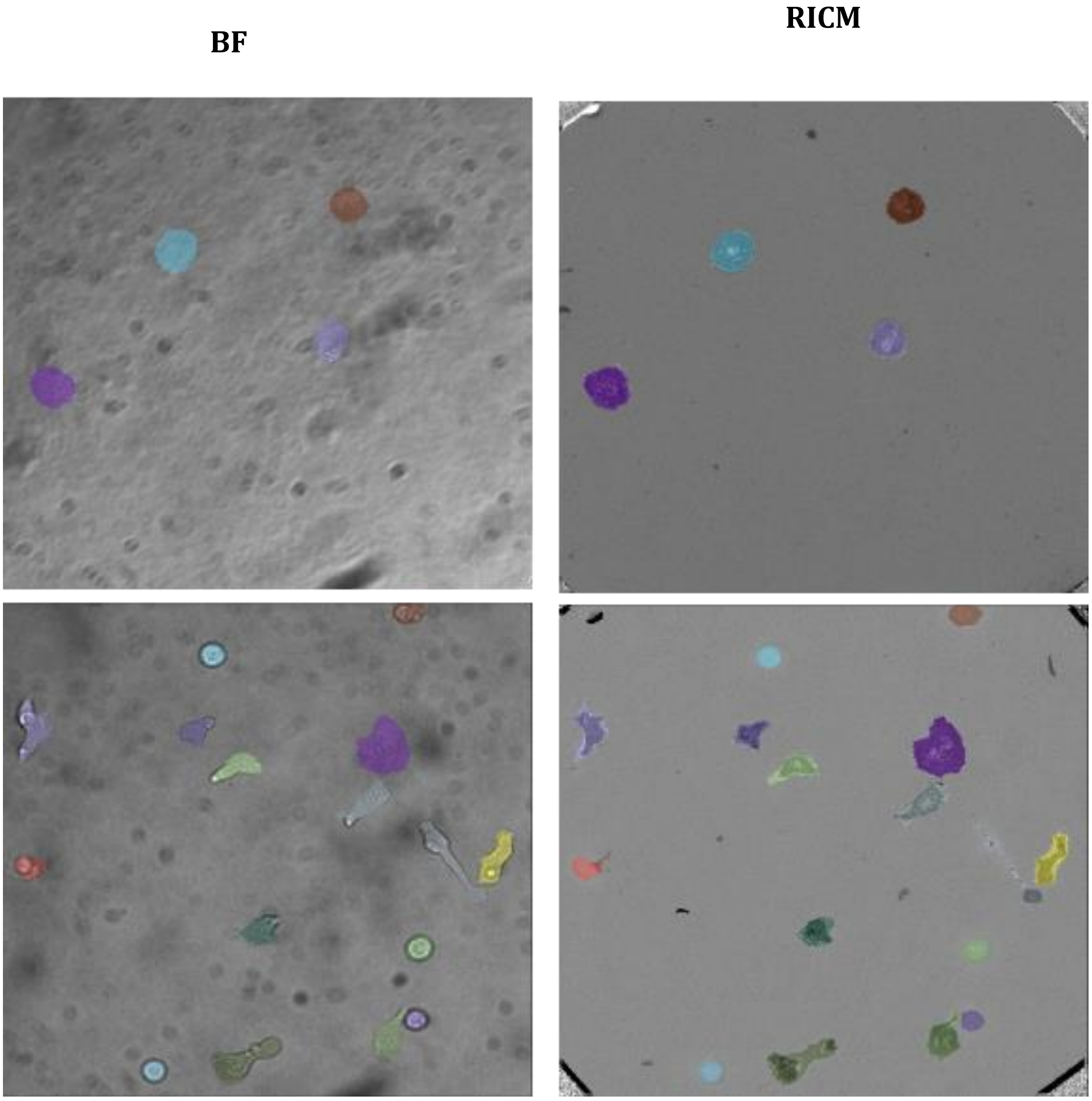


**Supplementary Figure 17. Cell tracking and trajectory analysis.** Cell tracking was performed using the trackpy module implemented within Celldetective. Segmented cells were linked across consecutive frames to reconstruct single-cell trajectories. This enabled time-resolved analysis of cell behavior, including quantification of adhesion area, cell motility, and detection of dynamic events such as the onset of spreading. Each trajectory was assigned a unique cell ID, allowing individual cells to be followed over time and enabling calculation of the fraction of cells undergoing specific events. Representative BF and RICM fields of view from two distinct conditions are shown: one on a surface without ICAM and one on a surface containing ICAM-1.

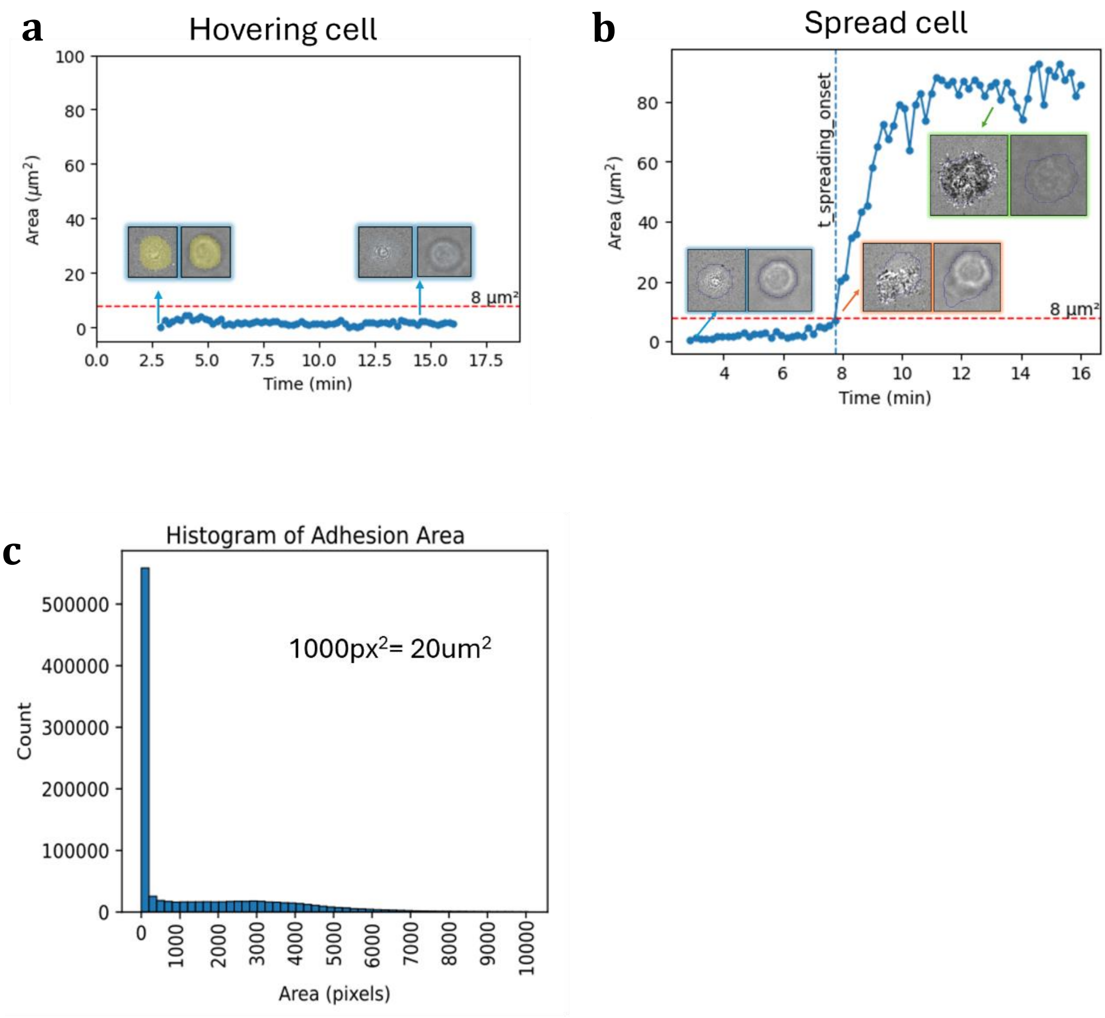


**Supplementary Figure 18. Determination of spreading onset.** Spreading onset was determined from the time evolution of the RICM close contact area, defined as the area within the segmented cell mask where the normalized RICM intensity was below 0.95. This metric captures the portion of the cell footprint in close contact with the surface. A threshold of 8 μm$^2$ was used to distinguish hovering from spreading cells: hovering cells remained below this threshold (as shown in (ii)), whereas spreading cells crossed it as their close-contact area increased (as shown in (ii)). The first time point at which the dark contact area exceeded 8 μm$^2$ was taken as the spreading onset time. As a consistency check, dark-contact-area values from all cells, all time points, and all conditions were pooled into a single histogram (as shown in (iii)). The pronounced peak below approximately 8 μm$^2$ supports the suitability of this threshold for separating hovering cells from the onset of spreading.

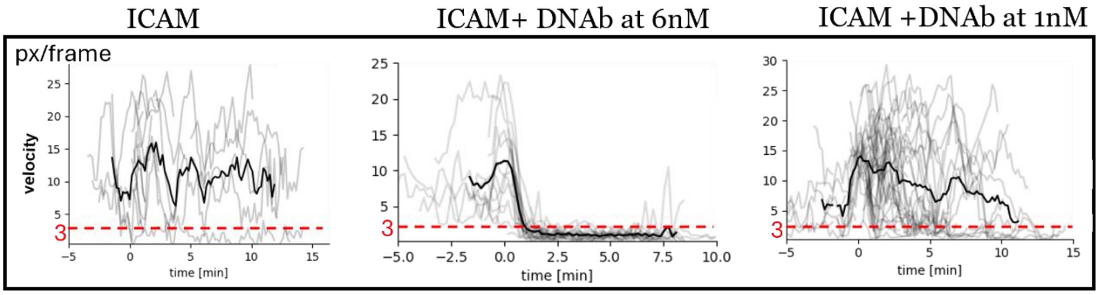


**Supplementary Figure 19.** Determination of arrest onset. Cell arrest was defined from the time evolution of cell velocity computed over a central window of 7 frames to reduce the noise. A threshold of 3 pixels per frame was used to distinguish motile from arrested cells: motile cells remained above this threshold as shown in case of ICAM, whereas cells undergoing arrest dropped below it as shown in case of ICAM and presence of high concentration of DNabs. The first time point at which the measured velocity fell below 3 pixels per frame (2um/min) was therefore defined as the arrest onset time. To assess whether this threshold was appropriate, velocity values were examined across cell tracks and conditions. The selected threshold was consistent with the transition from sustained motility to near-stationary behavior.

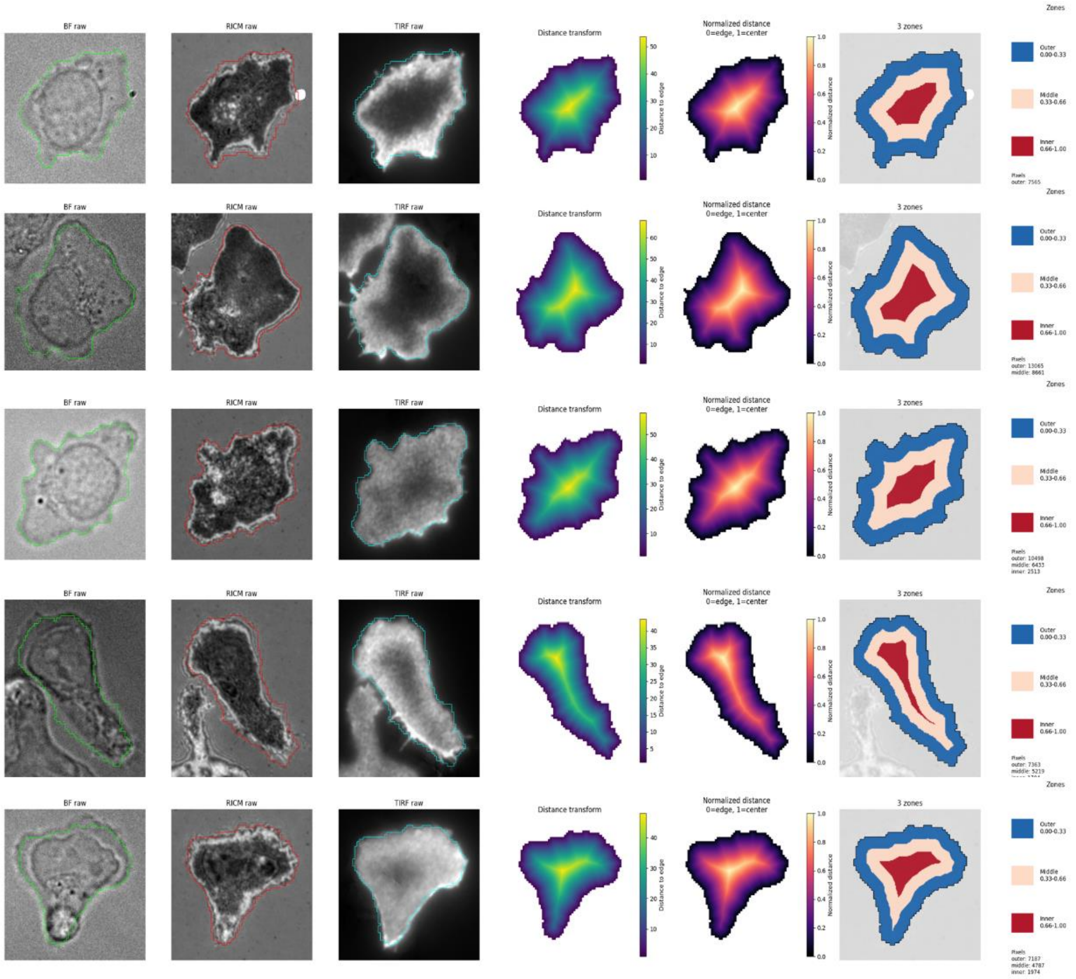


**Supplementary Figure 20. Quantification of CD45 exclusion.** Representative BF, RICM and TIRF images are shown for cells on ICAM-1 functionalized surface in the presence of DNabs (6 nM) as shown in i) or in the absence of DNabs (ii). Cell segmentation was was performed on the TIRF channel using the cellpose cyto3 model. A normalized distance transform was computed within each cell mask to divide the cell footprint into three radial zones defined as inner, middle and outer. The mean intensity in different zone is computed and ratio between inner and outer zone is computed and defined as [CD45 exclusion]$^{-1}$. Values close to 1 indicate little or no exclusion of CD45 from the cell center, whereas values below 1 indicate central exclusion of CD45.

| CD3 | $EC_{50}$ (nM) |
|---|---|
| 20 | 18.5 ± 8.4 |
| 30 cis | 24.8 ± 17.2 |
| 30 | 15.6 ± 0.6 |
| 40 | 10.9 ± 6.5 |
| Cn | 13.8 ± 6.5 |

| HER2 | $EC_{50}$ (nM) |
|---|---|
| 20 | 3.2 ± 0.8 |
| 30 cis | 5.3 ± 4.1 |
| 30 | 2.8 ± 0.5 |
| 40 | 2.8 ± 0.8 |
| nH | 2.8 ± 0.8 |

**Supplementary Table 1. Binding parameters of DNabs determined by flow cytometry.** $EC_{50}$ values were calculated from binding curves obtained, for CD3, on primary T lymphocytes from healthy donors and, for HER2, on MCF7-$HER2^+$ cell line. Data are reported as mean ± standard deviation (SD) from at least two independent experiments ($n \geq 2$). Abbreviation: $EC_{50}$, half-maximal effective concentration.

| | Date | | | |
|---|---|---|---|---|
| | 13/06/2025 | 04/07/2025 | 17/07/2025 | 29/07/2025 |
| MCF-7-HER2$^+$ + T cells | 0.93 | 0.91 | 0.78 | 0.92 |
| 20 | 0.78 | 0.39 | 0.26 | 0.74 |
| 30cis | 0.47 | 0.47 | 0.27 | 0.76 |
| 30 | 0.70 | 0.55 | 0.23 | 0.80 |
| 40 | 0.70 | 0.45 | 0.30 | 0.75 |
| Nef-CD3 | 0.92 | 0.81 | 0.81 | 0.93 |
| HER2-Nef | 0.92 | 0.83 | 0.86 | 0.91 |

**Supplementary Table 2. Survival of MCF-7-HER2$^+$ target cells after 4 h of DNab-mediated cytotoxicity.** The table reports, the fraction of surviving target cells (MCF-7-HER2$^+$) after 4 h for the highest DNab concentration (50 nM), quantified by live-cell imaging of propidium iodide (PI) uptake in Hoechst-stained target cell nuclei.

| Cell line | HER2 Expression (Receptors/cell) | EGFR Expression (Receptors/cell) |
|---|---|---|
| MCF-7 | $1.0x10^4$ | $1.2x10^3$ |
| MCF-7-HER2+ | $7.4x10^5$ | $0.7x10^3$ |
| A431 | $5.4x10^3$ | $2.8x10^5$ |

**Supplementary Table 3. Quantification of HER2 and EGFR surface expression on MCF-7, MCF-7-HER2$^+$, and A431 cell lines.**The table reports the number of HER2 and EGFR receptors per cell, on each cell line. These data were determined by flow cytometry using mouse (HER2) and human (EGFR) IgG calibration kits.

### Supplementary Videos

#### Supplementary Video 1 | T cell engagement on HER2-coated surfaces in the presence of DNAb

Time-lapse video showing T cell engagement on HER2-coated surfaces without ICAM in the presence of 6 nM DNAbs. Four image stacks corresponding to 30cis, 20, 30 and 40 DNAb engagers were combined horizontally using Fiji to compare spreading dynamics. Differences in engagement area, fraction of engaged cells and engagement rate can be observed across the different DNAbs. Time is shown as min. Scale bar, 20 µm.

#### Supplementary Video 2 | T cell arrest on HER2- and ICAM-coated surfaces in different concentration of DNAb

Time-lapse video showing T cell motility and arrest on surfaces coated with HER2 and ICAM in the presence of increasing DNAb concentrations. Four image stacks corresponding to 0 nM, 0.6 nM, 1 nM and 6 nM DNAb were combined horizontally using Fiji to compare arrest dynamics. The number of arrested cells increases with DNAb concentration, and cells arrest more rapidly at the highest concentration. Time is shown as sec. Scale bar, 20 µm.

#### Supplementary Video 3 | T cell-mediated killing of MCF-7-HER2$^+$ target cells in the presence of DNAb

Time-lapse video showing co-cultures of MCF-7-HER2$^+$ target cells and primary T cells at an effector:target ratio of 5:1 in the presence of DNAbs. Four conditions corresponding to 30cis, 20, 30 and 40 DNAbs were combined for comparison. Brightfield, Hoechst-labeled target nuclei, CFSE-labeled effector T cells and propidium iodide staining were acquired every 15 min over 5 h to visualize target-cell engagement and lysis. Time is shown in minutes. Scale bar, 40 µm.
